# Monocyte Dysregulation Defines an MDD-Specific Transcriptional Signature Closely Linked to Clinical MDD Traits

**DOI:** 10.1101/2025.08.12.669833

**Authors:** Laura Meier, Medhanie Mulaw, Verena Durner, Luca Schäfer, Carlos Schönfeldt-Lecuona, Verena Bopp, Heiko Graf, Markus Otto, Petra Steinacker, Henning Großkopf, Karin M. Danzer

## Abstract

**Objective:** Major depressive disorder (MDD) is a highly heterogeneous psychiatric condition contributing significantly to the global disease burden. In a subset of MDD patients inflammatory processes accompany the disease and this has also been reported for other psychiatric conditions like schizophrenia (SZ). However, little is known regarding immune deregulation in relation to clinical symptoms.

**Methods:** Cellular indexing of transcriptomes and epitopes by sequencing (CITE-Seq) of peripheral blood mononuclear cells (PBMCs) from patients with MDD (n=22), SZ (n=10), and healthy controls (HCs, n=26) was performed, yielding 633,284 high-quality single-cell profiles. Differential gene expression, cell composition, pathway enrichment, and weighted gene co-expression network analyses were conducted. MDD symptom severity was assessed using the Beck Depression Inventory-II (BDI-II), enabling the correlation of gene expression with individual disease symptoms.

**Results:** We identified disease-specific transcriptomic signatures for MDD as well as for SZ and shared gene expression changes between the two conditions. For MDD, monocyte numbers were elevated concomitant with a strong enrichment of interferon signaling. The MDD-specific signature involving all peripheral cell types correlated strongly with key symptoms of depression like sadness or pessimism. Reactome analysis identified nuclear factor kappa-light-chain-enhancer of activated B cells (NF-κB)–associated pathways among the genes positively correlating with disease symptoms and traits.

**Conclusion:** Our findings might help to stratify psychiatric patients and provide a multidimensional framework for investigating immune contributions to psychiatric disorders and highlight potential targets for therapeutic intervention. Clinical symptoms might be reflected on a molecular level paving the way for more targeted and effective interventions.

## Introduction

In their latest estimation, the WHO ranked major depressive disorder (MDD) as the single largest contributor to global disability and suicide as one of the leading causes of death worldwide ^1,2^. While there are many theories trying to explain the pathophysiology of MDD, the disease mechanisms remain poorly understood and are highly heterogenous. Genetic susceptibility can contribute to developing depression; other triggers are for instance chronic stress and traumatic experiences ^3^. Emerging evidence suggests that a subset of patients suffering from depression shows a pattern of inflammation, indicated by elevated cytokine and acute-phase protein levels ^4,5^. Patients with an autoimmune disorder have an increased risk of developing MDD and anti-inflammatory treatment has been reported to have beneficial effects on depressive symptoms ^6–8^. Additionally, inflammation may contribute to the treatment unresponsiveness that concerns a significant number of patients with MDD ^9^. Supporting evidence for a role of inflammation in depression comes from a study reporting that endotoxin administration induces depressed mood and reduced striatal activity as a measure for anhedonia in study participants ^10^. Social withdrawal, motivational deficits and fatigue – the so called “sickness behavior” that shares many symptoms with MDD – are evolutionary purposeful reactions to infection and disease ^11,12^. While inflammation was mainly driven by injuries or pathogen contacts in earlier times, nowadays diseases of affluence like diabetes, obesity or metabolic syndrome contribute to a general pro-inflammatory tendency and thereby potentially to rising depression diagnosis ^13^. Growing evidence suggests that inflammation-reducing treatments may have antidepressant effects in a subset of individuals with major depressive disorder (MDD). However, further research is needed to identify which patients are most likely to benefit from this approach ^14,15^.

Evidence of immune system involvement in MDD also extends to schizophrenia (SZ). For example, prenatal exposure to infections such as Herpes simplex virus (HSV) or influenza has been associated with an elevated risk of developing SZ in adulthood. Furthermore, as seen in MDD, changes in cytokine levels have been reported in individuals with SZ, highlighting inflammation as a potential common pathway in the pathophysiology of both disorders. ^16,17^ The precise mechanisms in which the peripheral immune system contributes to psychiatric diseases remain unclear. An *in vitro* stimulation of peripheral blood mononuclear cells (PBMCs) in MDD patients with lipopolysaccharide showed an increased immune cell response that is associated with anhedonia ^18^. Even though the brain has been considered immunologically privileged for a long time, it is certainly not shielded from inflammatory influences from the periphery. For instance, the blood-brain barrier (BBB) contains transporters to selectively convey cytokines, including interleukin (IL)-6 and tumor necrosis factor (TNF), from one side to the other ^19–21^. While the BBB tightly regulates trespassing of molecules and cells under physiological conditions, its integrity suffers distinctly during inflammation, allowing an influx of cytokines and immune cells from the periphery ^22^. Indeed, a recent study demonstrated that chronic social defeat stress (CSDS) in mice elevates circulating Ly6C^hi^ monocytes and neutrophils, enabling their migration across the BBB into the brain. Moreover, single-cell transcriptomics revealed upregulation of matrix metalloproteinase 8 (Mmp8) in brain Ly6C^hi^ monocytes, which induces extracellular space alterations in the nucleus accumbens, contributing to social withdrawal behavior in CSDS mice. A principal elevation of monocytes and neutrophils in MDD patients could also be confirmed ^23^.

Further understanding of the underlying mechanisms that connect peripheral inflammation to psychiatric disorders contributes significantly to the identification of eligible treatment options. In this study, we utilized cellular indexing of transcriptomes and epitopes by sequencing (CITE-Seq) ^24^ to analyze single-cell gene expression and cell surface protein profiles of PBMCs from patients with MDD and SZ. By integrating clinical patient data with this multidimensional dataset, we provide a unique framework to explore potential links between molecular profiles and clinical outcomes.

## Methods and Materials

### Study participants and ethical approval

Human blood sample collection was approved by the Ethic Committee of Ulm University (proposal number 405/19, year 2019) and all volunteers signed informed written consent prior to participation. MDD and SZ patients were assessed and diagnosed by board-certified psychiatrists based on the DSM-5 criteria. Symptom severity was evaluated using the 21-itemed Beck Depression Inventory (BDI) – II for patients with MDD or the positive and negative scale (PANSS) for patients with SZ. Comprehensive medical history and detailed biographical data were obtained for all participants. Patients with acute or chronic inflammatory diseases were excluded from participation. Healthy controls without neurological, psychiatric or inflammatory conditions were recruited to match age and sex of the patient cohort. 15-37·5 mL venous blood were drawn and collected in a Sarstedt ethylenediaminetetraacetic acid (EDTA) Monovette®.

### PBMC isolation

PBMCs were isolated from venous blood samples by density gradient centrifugation with Histopaque®-1077. After cautiously transferring the PBMCs, they were washed two times with Dulbecco’s phosphate buffered saline (DPBS) before resuspending them in Cell Staining Buffer (BioLegend®) and filtering through 70 µm and afterwards 40 µm Flowmi® Cell Strainer (Merck).

### Antibody labelling of PBMCs

To reduce unspecific binding 500,000 PBMCs were treated with Human TruStain FcX™ Fc receptor blocking solution (BioLegend®) for 10 min at 4 °C. Afterwards the cells were labelled with the TotalSeq™-B Human Universal Cocktail, V1.0 (BioLegend®) for 30 min at 4 °C. Cells were washed 3 times with Cell Staining Buffer (BioLegend®) before resuspending them in an appropriate volume of the same buffer.

### 3’ Gene expression and antibody derived tag library preparation

The library preparation was conducted following the instructions provided by 10x Genomics (CG000317 Rev C). Using the Single Cell 3’ version 3.1 (Dual Index), 20,000 cells were loaded for a recovery of around 12,000 cells. cDNA was amplified for 11 cycles and afterwards size selected using SPRIselect™ beads (Beckman Coulter™) for separation of the amplification product for 3’ RNA and antibody derived tag (ADT) library construction. Amplification cycles for the gene expression library were adjusted depending on the cDNA concentration. For the ADT library 10 cycles of amplification were performed.

### Sequencing

Next generation sequencing was performed by Novogene using the NovaSeq 6000 (Illumina®).

### Data Analysis

Raw single-cell data was demultiplexed using Cell Ranger ^25^. RNA and ADT data quality control, normalization, and integration were performed using the R package Seurat ^26–30^. Additional downstream analyses were performed using additional R ^31^. RNA was normalized using the sctransform ^32^ by regressing out mitochondria content and cell cycle phase score. ADT data was normalized using Centered Log Ratio (CLR) followed by data scaling. To adjust for batch effect, data integration was performed at the samples level using Principal Component (PC) embeddings-based harmonization. To adjust for batch effect and data integration, samples were first merged (ensuring a union list of features across all samples with unique cell IDs). To integrate the RNA data, we performed PCA on the SCT slot, followed by data Principal Component (PC) embeddings-based harmonization with k-means initialization set to 100 and a maximum iteration of 50. Convergence plots evaluated data harmonization. For ADT, PCA was performed on the scaled merged data, followed by the harmonization steps as discussed for the RNA dataset. UMAP and FindNeighbors analysis (Seurat) was then performed (for both RNA and ADT) by using the Harmony data reduction slot.

## Results

Transcriptomic signatures of MDD and SZ reveal shared pathways and overlapping affected cell populations We generated a multi-modal, multi-cohort CITE-Seq dataset of PBMCs from 22 MDD patients, 10 SZ patients and 26 HCs profiling both gene expression and protein abundance simultaneously in individual cells (figure 1 A, supplementary table 1). MDD and SZ were diagnosed based on standardized criteria (see methods). By utilizing the 10X Genomics single-cell mRNA sequencing technique alongside antibody-derived tags (ADTs), for protein detection, we acquired a total of 633,284 high-quality single cells with simultaneous transcriptome and epitope profiles, comprising 224,227 cells from MDD patients, 119,257 cells from SZ patients and 289,800 cells from HCs. We then integrated all samples into a fully merged, integrated dataset by carefully modelling the batch effects with Principal Component (PC) embeddings-based harmonization, aiming for a robust clustering of cell communities independent from batch and the donor disease status. To determine cell identity and group cells based on transcriptome profiles and ADT surface marker expression programs, we leveraged both RNA and ADT modalities by employing the weighted-nearest-neighbor (WNN) framework, resulting in 31 cell clusters: T cells (naïve, proliferating, effector memory (TEM) and central memory (TCM) CD4 T cells; cytotoxic CD4 T cells (CD4 CTL); naïve, proliferating CD8, CD8 TEM and TCM cells, regulatory T cells (T_reg_); double negative T cells (dnT); γδ T cells (gdT); mucosal-associated invariant T cells (MAIT)), B cells (naïve, intermediate and memory B cells; plasmablasts), NK cells (proliferating and CD58^bright^ NK cells), monocytes (CD14 and CD16 monocytes), dendritic cells (type 1 and type 2 conventional dendritic cells (cDC1/2); A XL^+^ SIGLEC6^+^ dendritic cells (ASDC); plasmacytoid dendritic cells (pDC)), innate lymphoid cells (ILCs), hematopoietic stem and progenitor cells (HPSCs), erythrocytes and platelets.

**Figure 1:**
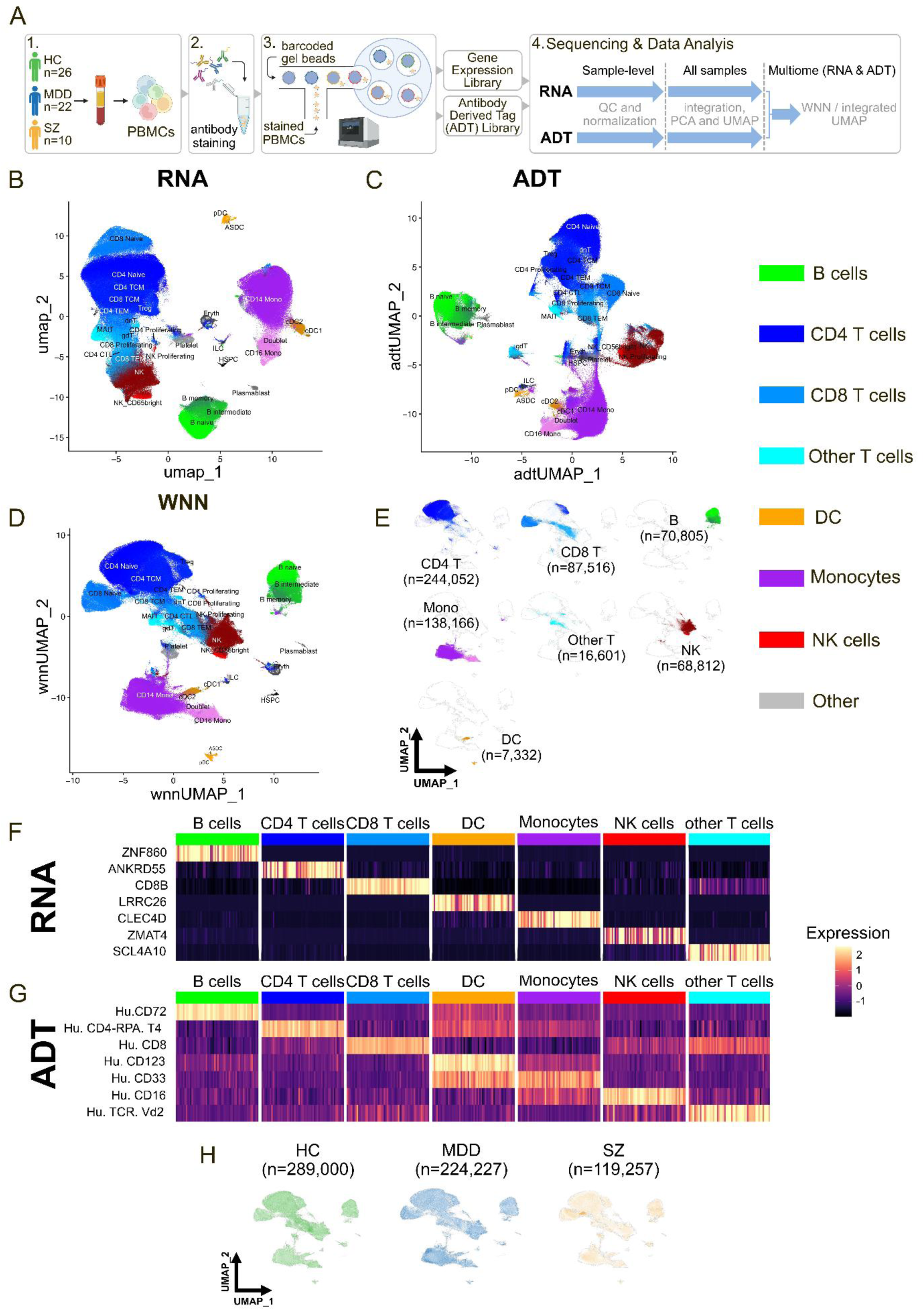
Characterization of 633,284 high-quality single cells. **A** Schematic representation of the CITE-Seq library preparation protocol. 1. EDTA-blood was drawn from 26 healthy participants, 22 MDD patients and 10 SZ patients. PBMCs were isolated using a density gradient. 2. PBMCs were stained with oligonucleotide-labelled antibodies targeting 134 unique cell surface proteins. 3. Using the microfluid-based approach from 10x Genomics, two distinct libraries can be prepared, reflecting the transcriptome and ADTs. 4. Sequencing was conducted at Novogene (Munich, Germany) using the Illumina NovaSeq 6000 and computational analysis started. After quality control (QC) according to current standards and normalization on a sample-level, data of all samples was integrated, and principal component analysis (PCA) is conducted, and two-dimensional uniform manifold approximation and projection (UMAP) plots are generated. In the last step, RNA and ADT derived data is integrated for a multiomic approach. **B-E** UMAP plots based on the single cell gene expression (RNA, **B**), antibody derived tags (ADT, **C**), and the integrated data based on the weighted nearest neighbor principle (WNN, **D**). **E** shows UMAPs of major cell types showing distinct clustering patterns based on multimodal information (gene expression and ADT). Detailed annotations revealed 31 different cell types. Representative top markers defining major cell types using gene expression (**F**) and ADT (**G**) datasets. **H** UMAPs split by patient group show a relative uniform representation of the different clusters (cell numbers by patient groups are indicated).

Cell clusters were classified into major immune cell types (B cells, CD4 T cells, CD8 T cells, other T cells, DCs, monocytes, NK cells) using established marker genes and ADTs (figure 1 F, G). Cells not falling into the aforementioned major cell types were defined as “other” ^27^ and comprise heterogenous cell types that cannot be analyzed as a group and hence were excluded from further analysis. Major cell types were evenly distributed across disease groups (MDD, SZ, HC), as well as sex and age groups, with no significant overrepresentation of subclusters in any condition (figure 1 H, supplementary figure 1).

We next sought to identify gene expression changes associated with MDD. We conducted single-cell differential gene expression analysis with MDD and healthy controls (HC) and identified 3,087 upregulated and 3,950 downregulated differentially expressed genes (DEGs), while pseudobulk analysis (aggregating all donor cells) showed 112 upregulated and 584 downregulated genes. Similarly, for PBMCs from SZ patients compared to controls, single-cell analysis revealed 2,094 upregulated and 4,527 downregulated DEGs, with pseudobulk confirming 122 upregulated and 99 downregulated genes. Notably, nearly all pseudobulk DEGs were also identified in single-cell analysis, highlighting a transcriptional core signature for each disease (figure 2 A and B, supplementary table 2).

**Figure 2:**
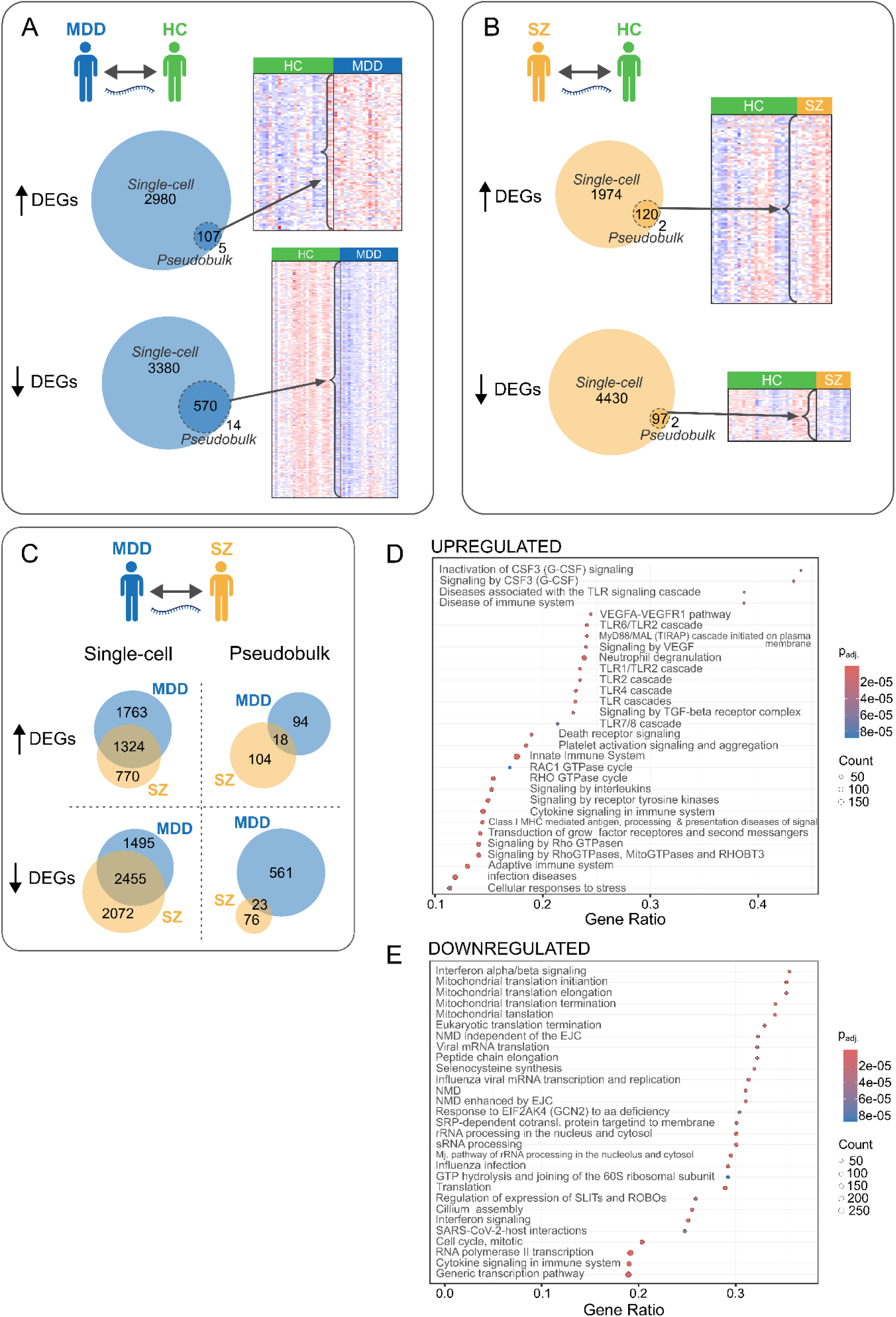
Disease-specific transcriptomic changes and shared disease mechanisms. Comparative differential gene expression analysis of a single-cell versus a pseudobulk approach in MDD and SZ. **A** Gene expression based differential analysis between MDD and HC (single-cell level) identified 3087 up- and 3950 downregulated genes. Pseudobulk analysis (where cell counts were pooled by patient groups and cell types; see methods for further details) yielded 112 up- and 584 downregulated genes. Both analysis methods share 107 up- and 570 downregulated DEGs. **B** In the SZ to HC comparison, the single-cell approach showed 2094 up- and 4527 downregulated genes. Pseudobulk analysis identified 122 up- and 99 downregulated genes. Both analysis methods share 120 up- and 97 downregulated DEGs. **C** A comparison of MDD and SZ differential expression analyses showed 1324 shared up- and 2455 shared downregulated genes from the single-cell differential expression analysis. Furthermore, there were 18 and 23 genes (up and downregulated, respectively) in common between MDD and SZ in the pseudobulk approach. **D + E** By taking the DEGs overlapping between MDD and SZ, we performed a Reactome pathway analysis. Dot plots are shown indicating the gene ratio, significance (p_adj_.) and gene counts. Toll-like receptor (TLR), VEGF, RhoGTPase, and Signaling by CSF-3 were amongst the upregulated gene-based pathways that showed significant overrepresentation and an FDR cutoff of 0.05. In the downregulated gene-based analysis, transcription related processes including Major pathway of rRNA processing in the nucleolus and cytosol, Nonsense mediated decay (NMD), and Translation showed significant enrichment. We additionally noted that Interferon and Cytokine signaling were statistically significantly overrepresented.

Comparing MDD and SZ revealed 1,324 shared upregulated and 2,455 shared downregulated DEGs in single-cell analysis (18 up- and 23 downregulated DEGs in pseudobulk), suggesting common disease pathways. Reactome pathway analysis was conducted on the single-cell DEGs overlapping between MDD and SZ to gain further mechanistic insight into commonly affected pathways contributing to psychiatric pathology. Among the upregulated pathways terms associated with CSF3-, TLR- or VEGF-signaling and interleukin and cytokine signaling (figure 2 D) were identified. Downregulated terms included interferon signaling, mitochondria-associated functions, translation and rRNA processing (figure 2 E).

### MDD disease signature converges on interferon signaling and is reflected in monocytes

Since monocyte numbers have been reported to be increased in MDD ^23,33^ we analyzed cell type composition for all major cell types (B cells, CD4 T cells, CD8 T cells, other T cells, monocytes, NK cells, DC cells and other cell types). We could indeed confirm elevated monocyte numbers in MDD compared to HC (*p_Holm-adj._* = 0·01, figure 3 A). While B cell counts were significantly higher in MDD compared to SZ (pHolm-adj. = 0·02, figure 3 A), other cell types showed no significant differences.

**Figure 3:**
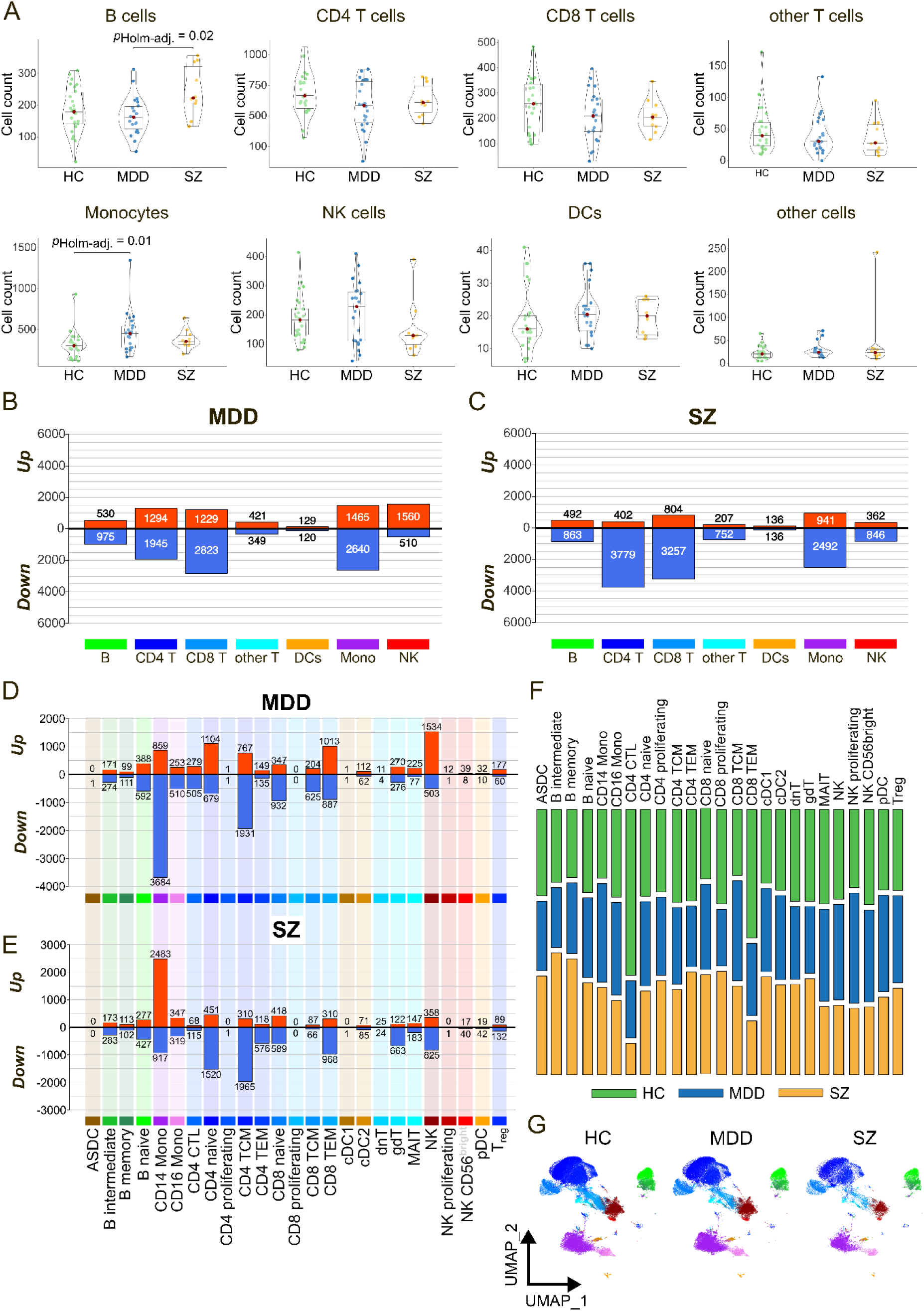
Differences in cell type composition and gene expression. **A** Comparison of cell numbers of all major cell types between HC, MDD and SZ using Kruskal-Wallis and Dunn testing. Significant differences were found in the B cell populations of MDD and SZ with a p_Holm-adj._ = 0·02 and in monocytes between HC and MDD with a p_Holm-adj._ = 0·01. **B + C** Numbers of differentially expressed genes (DEGs) (padj. = 0·05) in each major cell type for MDD compared to HC (**B**) and for SZ compared to HC (**C**). **D + E** Numbers of DEGs (p_adj._ = 0·05) per cell subcluster (detailed) in the MDD group (**D**) and the SZ group (**E**) in comparison to HC. **F** Cell number contribution per disease group (green=HC, blue=MDD, yellow=SZ) in each subclusters shows no significant overrepresentation. **G** WNN-UMAPs show the distribution of all cellular subclusters for each study group and all cell types are equally represented.

To evaluate their individual contributions differential expression events were analyzed across the seven major cell types. Increased resolution enhanced the detection of DEGs, revealing specific transcriptomic alterations in major cell types (figure 3 B). Transcriptomic changes were prominently observed in CD4 T cells, CD8 T cells and monocytes in both MDD and SZ. Notably, DEGs were predominantly downregulated in both disease groups. In MDD, NK cells are the only cell type, where upregulated DEGs predominate.

To gain a deeper insight into transcriptional changes, DEGs were determined for all 25 detailed cell clusters. No significant differences were apparent in the detailed subclusters regarding cell number contribution from the MDD, SZ and HC group (figure 3 F) and all subclusters were well represented in the groups (figure 3 G). CD14 monocytes exhibited the highest number of DEGs in both MDD and SZ, with distinct expression patterns: in MDD, most DEGs were downregulated, while in SZ, upregulated genes predominated, suggesting divergent disease mechanisms in this cell type.

Gene set enrichment analysis (GSEA) revealed that MDD signature converges in interferon (IFN) signaling and is mainly present in monocytes. While also some other cell types show some enrichment in IFN signaling (supplementary figure 2), highly significant enrichment was detected in both CD14 and CD16 monocytes (figure 4). The same trend was observed for IFN response genes, where significant enrichment specifically for target genes of type I and II IFN was ascertained (figure 4).

**Figure 4:**
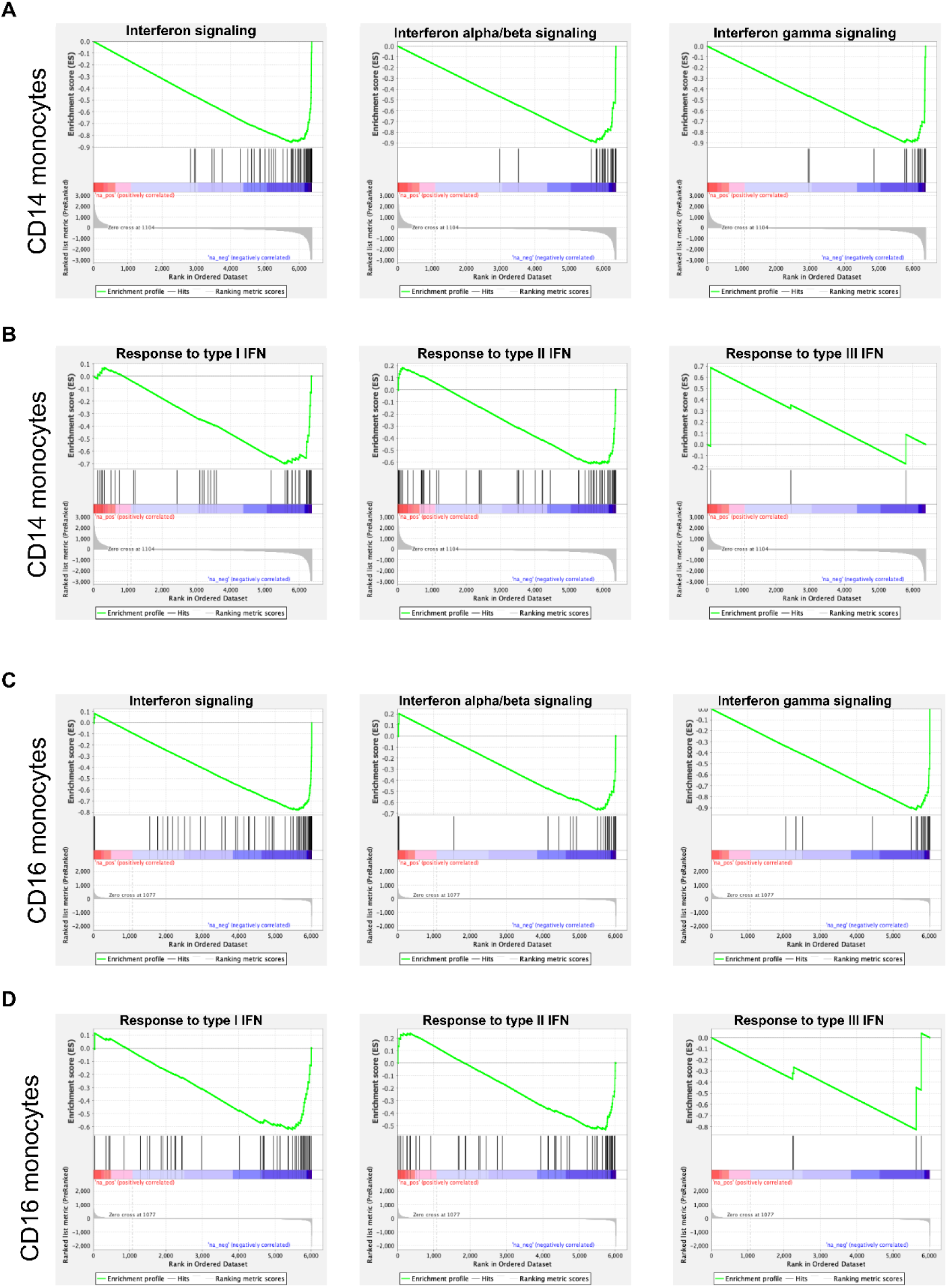
Enrichment plots of IFN signaling and IFN response genes for CD14 and CD16 monocytes. GSEA shows a highly significant enrichment of gene sets of IFN signaling, IFN alpha/beta signaling and IFN gamma signaling in CD14 (**A**) and CD16 monocytes (**C**) from MDD patients compared to HC. Significant enrichment was also present for IFN response genes, specifically genes responsive to type I and II IFN in CD14 (**B**) and CD16 monocytes (**D**).

A highly significant enrichment of IFN signaling was also identified in CD14 and CD16 monocytes in SZ, suggesting shared disease mechanisms in these cells (supplementary figure 3). Further, we found the MDD core signature to be represented across various cell types in SZ (supplementary figure 4) which substantiates the idea of common psychiatric disease mechanisms.

### Sadness and pessimism present the strongest association with MDD transcriptomic signature

To identify clinical traits that relate with the MDD disease signature, clinical symptoms were ranked based on association to MDD gene expression changes. The symptoms of sadness, pessimism and loss of interest presented the strongest association (figure 5 A). To examine the relation of gene expression changes in MDD and clinical symptoms, we performed a weighted gene co-expression network analysis (WGCNA), clustering the 677 genes from the MDD core signature (common MDD gene signatures between single-cell and pseudobulk) into 5 modules of similar expression pattern across all MDD patients (modules “blue, brown”, etc.). Correlation of the module expression scores with clinical parameters revealed that the expression of the brown module was strongly positively correlated to the depression disease score (BDI-II) and sadness, while age and BMI did not correlate with gene modules (figure 5 B).

**Figure 5:**
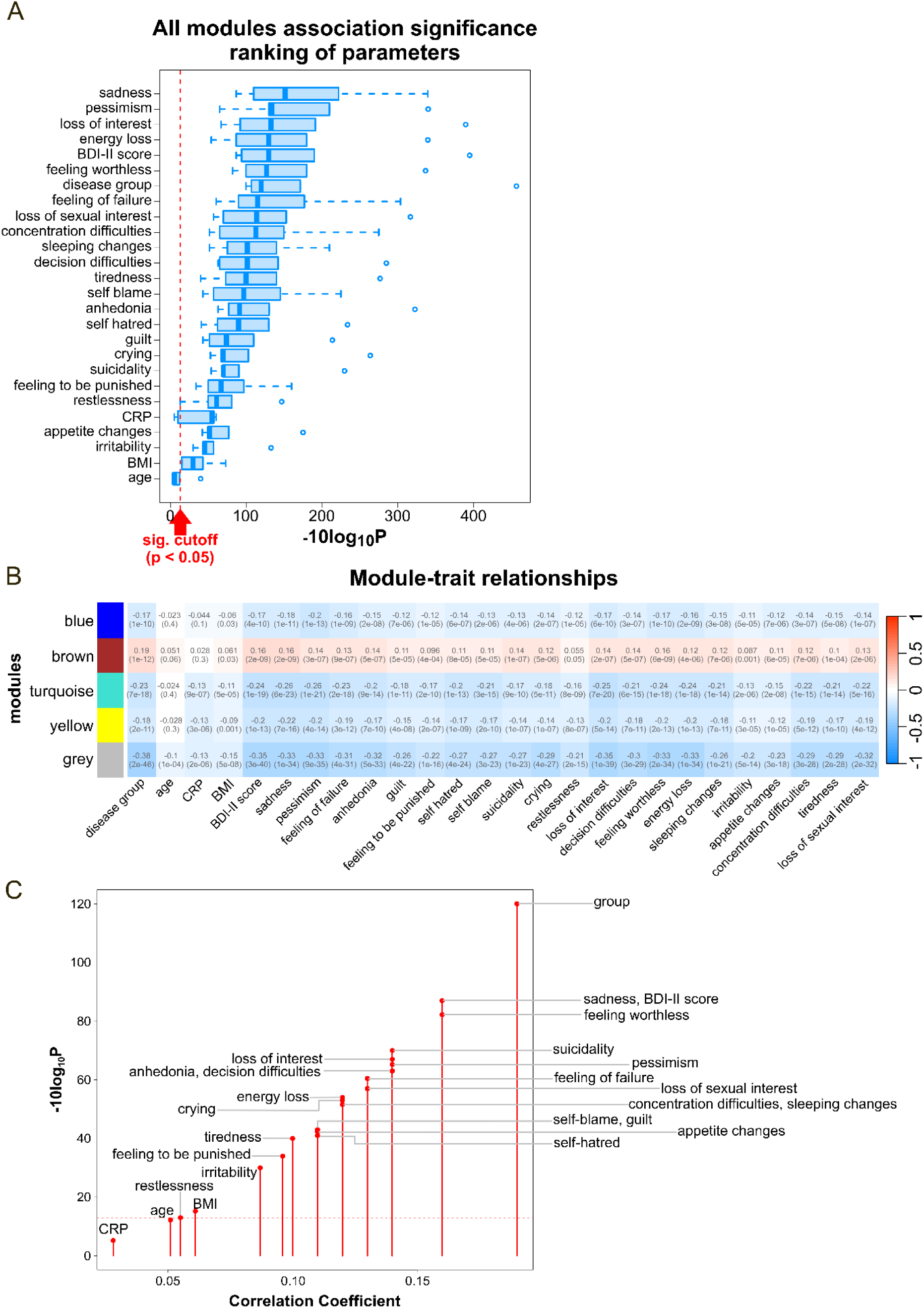
Differential gene expression and the association with symptoms and traits. **A** By subsetting the DEGs in the transcriptomic MDD core signature, generated by overlapping single-cell and pseudobulk analysis for the MDD to HC comparison (figure 2 A), we identified 5 groups (modules) of highly correlated genes. The association of each module to clinical symptoms and traits are shown, ranked by the median of the significance across all modules (with a cutoff of p < 0·05; x axis shows −10log_10_P transformed p-values). Sadness, pessimism, and loss of interest ranked highest while age showed the lowest overall association to the identified modules. **B** The 5 identified modules (“blue”, “brown”, “turquoise”, “yellow” and “grey”) and their correlation with different parameters are displayed in a heat-map. Overall, the modules had significant negative correlation to the parameters, where only module ‘brown’ shows a positive correlation. **C** Correlation coefficients of the distinct parameters with module “brown”. Highest correlation is found for the different disease groups followed by symptoms of sadness and the feeling of worthlessness as well as the overall BDI-II score. The weakest correlation is found for CRP values, the age, the symptom of restlessness and the body mass index (BMI).

To gain further mechanistic insight, we analyzed the functional enrichment of Reactome terms of the genes in the brown module. We found that genes involved “NF-κB is activated and signals survival”, “p75NTR signals via NF-κB” and “N-glycan trimming in the ER and Calnexin/Calreticulin cycle” were significantly enriched (**p < 0·01, q > 0·05) in module brown.

To find the clinical parameter that was most affected by gene expression changes in the brown module the correlation coefficients were ranked. Here, sadness, the BDI-II score and feeling worthless showed the strongest correlation (figure 5 C). These results suggest that sadness and other psychiatric symptoms can be reflected in a transcriptomic fingerprint in PBMCs.

## Discussion

In this study, we performed a multimodal single-cell analysis of the transcriptome and epitome of PBMCs from MDD and SZ patients in comparison to HC. A total of 633,284 high-quality single cells were examined and characterized. We unveiled a disease-specific transcriptomic core signature for MDD and SZ each, as well as shared gene expression changes in both mentioned diseases. Like previous studies, we could confirm an overrepresentation of monocytes in MDD ^23,33^. Moreover, we identified CD14 monocytes as a key cell type harboring an MDD specific transcriptional signature and a strong enrichment of IFN signaling in monocytes. For the first time in MDD, transcriptomic changes on a single cell level were correlated with disease parameters and symptoms depict a strong correlation for the symptoms of “sadness” and “pessimism” with a transcriptomic signature.

While bulk RNA sequencing of PBMCs from MDD patients with different treatment responsiveness could not detect transcriptional changes ^34^, we could provide evidence for disease-specific transcriptomic signatures in our MDD or SZ cohorts using single-cell differential gene expression analysis and could verify those in pseudobulk analysis. Shared DEGs between the single-cell and pseudobulk approach hereby describe a transcriptomic disease core signature for each disease. In MDD versus HC there are 107 upregulated disease core transcripts including genes like *NFKBIA*, *CD55* and *TNFAIP3* that are involved in inflammatory processes. While these genes have not yet been implicated in classical psychiatric conditions, there is evidence of their contribution to neurological diseases like Alzheimer’s ^35^ or neuropsychiatric systemic lupus erythematosus ^36^. Upregulation of *CD55* has been reported during autoimmune inflammatory processes in the CNS and is suggested to play a role in neuroprotective processes ^37^. Its increased expression in MDD might have compensatory effects in the pro-inflammatory state that affects at least a subset of patients ^38^. 570 genes are downregulated in the disease core signature. Among them, the transcription factor *SP4* has already been implicated in other psychiatric diseases and a reduced expression has also been reported in PBMC from first-time psychosis patients ^39,40^.

We found a highly significant enrichment of IFN signaling in CD14 and CD16 monocytes in MDD. Type I IFNs like IFN-alpha and -beta are components of the innate immune system and are necessary for fighting infections and limiting cell growth ^41–43^. Type II IFN, IFN gamma, is involved in adaptive and innate immune responses and antitumor processes ^44^. Previously, elevated expression levels of genes associated with IFN alpha/beta signaling in patients with recurrent MDD using whole blood RNA sequencing have been reported ^45^. In line with this, IFN-alpha administration, used to treat hepatitis C virus infection or malignancies, induced depression in a significant number of patients ^46,47^. IFN-alpha-associated CNS inflammatory responses have been suggested to interact with monoamine (serotonin) metabolism, which is associated with depression ^48^. In contrast to these studies, we found a downregulation of key players of INF signaling (*JAK1*, *JAK2* and *STAT1*) in CD14 and CD16 monocytes of MDD patients. It might well be that this points to compensatory mechanisms of certain IFN-associated genes in an otherwise elevated pro-inflammatory state. Supporting evidence for this hypothesis comes from a single nuclei transcriptomic study, that showed downregulation of IFN signaling in microglia in female MDD patients using post-mortem prefrontal cortex samples ^49^. Also Morrison *et al.* demonstrated in prefrontal cortex postmortem material from MDD patients a reduced *IL1A* gene expression ^50^.

In summary, transcriptomic changes in MDD are reportedly diverse ^34,45,51^ which reflects the heterogeneity in the disease itself. Previous studies have shown that peripheral and brain-specific gene expression can correlate ^52^, suggesting that integrating peripheral transcriptomic data with single-cell transcriptomic data from post-mortem brain samples of MDD patients could provide a more comprehensive picture of the involvement of INF signaling in MDD. Apart from differential gene expression, the multimodal single-cell and ADT analysis allows precise annotation of the cells’ identity and composition. A single-cell RNA and ATAC (assay for transposase-accessible chromatin) sequencing study revealed differences in numbers and transcription profiles of CD4 naïve T cells ^53^. In contrast to Sun *et al.* ^53^, we found a significant increase in monocytes in MDD compared to HC. This finding in MDD patients has previously been recapitulated in a CSDS mouse model, where an increase of Ly6C^hi^ inflammatory monocytes and a decrease of B cells has been observed ^23^. Further, a study with 22 MDD patients and 14 HCs reported an overrepresentation of intermediate monocytes while classical monocyte frequency was decreased ^33^. While our transcriptional approach confirmed an increase in total monocyte counts, no significant differences were observed in monocytic CD14 and CD16 subtypes. Further evidence for the strong involvement of monocytes in depression comes from our MDD transcriptomic signature. Here monocytes are strongly overrepresented and contribute distinctly to differential gene expression with 3,864 downregulated genes in MDD. In contrast, SZ monocytes showed an opposing direction of differential regulation in CD14 monocytes which might indicate that the same cell type is driving the diseases’ transcriptional manifestation, however, depending on the transcriptional characteristics, the disease manifestation towards MDD or SZ differs.

Accumulating evidence for genetic susceptibility for different psychiatric diseases like MDD, SZ, bipolar disorder, autism spectrum disorder or attention-deficit/hyperactivity disorder exists ^54,55^. However, from the viewpoint of precision medicine, the identification of underlying mechanisms for traits and individual symptoms is highly relevant. Since treatment resistance is a major problem in MDD patients, addressing individual symptoms instead of the diagnosis and stratification of patients with targeted therapies could ameliorate disease burden significantly. This study might help to associate transcriptomic signatures with specific symptoms of mood disorders (like sadness or anhedonia), which has never been examined this closely before. Ward *et al.* investigated the genetic background of the rather generic symptom of “mood instability” and identified 46 loci, 244 significant genes and 6 gene sets associated with the trait ^56^. We utilized the BDI-II questionnaire to assess the correlation between individual symptom severeness and traits like age and BMI and transcriptional changes in the disease signature. The strongest association is shown for the symptoms of “sadness” and “pessimism”, which are key features of depression, experienced by most patients. Interestingly, the association of traits like age, BMI and CRP is rather low, indicating an overestimated impact of these factors on gene expression changes compared to disease symptoms of MDD. By refining the analysis and clustering genes from the transcriptomic core signature with similar expression patterns, gene module “brown” has been identified to have a positive correlation to MDD symptoms. The brown module has the strongest correlation of transcriptomic changes for the symptoms of sadness and feeling of worthlessness. Reactome pathway analysis of the brown module detected an enrichment in the terms “NF-κB is activated and signals survival”, “p75NTR signals via NF-κB”, and “N-glycan trimming in the ER and Calnexin/Calreticulin cycle”. Two of the 3 reactome terms involve NF-κB signaling, which is crucial in inflammatory responses and regulations and further substantiates the association of inflammation and depression ^57^. Indeed, evidence that NF-κB pathway activation is involved in the pathophysiology of depression and its downregulation might have antidepressive effects has been suggested previously ^58,59^. Ward *et al.* reported association of depression with complex 1 associated genes (*NDUFAF3, NDUFS3* or *MTCH2*) involved in energy production by mitochondria ^56^. Similarly, we could observe a significant downregulation of mitochondrial components (like *NDUFB3* and *NDUFAF1)*. However, they are not positively correlated with specific disease traits.

In conclusion, we could provide evidence for transcriptional and cell composition changes in MDD compared to HC in a multimodal, high-quality single cell data set. Further, we identified a disease-specific transcriptional core disease signature for MDD and SZ, as well as some shared transcriptomic profiles. In MDD, monocytes are distinctly contributing to the transcriptomic changes and a strong enrichment for IFN signaling was identified in CD14 and CD16 monocytes. Further we could show that symptom manifestation can be reflected in gene expression changes.

## Acknowledgments

We thank all probands for their participation in this study. This work was funded by the Boehringer Ingelheim Ulm University Biocenter (BIU), the Deutsche Forschungsgemeinschaft (DFG) 637 Emmy Noether Research Group DA 1657/2-1, and SFB 1506 638 (Aging at Interfaces).

## Abbreviations

ADT: antibody derived tag
ASDC: AXL^+^ SIGLEC6^+^ dendritic cell
BBB: blood brain barrier
BMI: body mass index
CD4 CTL: cytotoxic CD4 T cell
cDC1/cDC2: type 1/type 2 conventional dendritic cell
CITE-seq: cellular indexing of transcriptomes and epitopes by sequencing
CLR: Centered Log Ratio
CRP: C-reactive protein
CSDS: chronic social defeat stress
DC: dendritic cell
DEG: differentially expressed gene
DFG: German Research Foundation (Deutsche Forschungsgemeinschaft)
dnT: double negative T cell
DPBS: dulbecco’s phosphate buffered saline
EDTA: ethylenediaminetetraacetic acid
gdT: γδ T cell
GSEA: Gene set enrichment analysis
FDR: false discovery rate
HC: healthy control
HPSC: hematopoietic stem and progenitor cell
IFN: interferon
IL: interleukin
ILC: innate lymphoid cell
MAIT: mucosal-associated invariant T cells
MDD: major depressive disorder
MMP8: matrix metalloproteinase 8
NF-κB: nuclear factor kappa-light-chain-enhancer of activated B cells
NK: natural killer cell
PBMC: peripheral blood mononuclear cell
pDC: plasmacytoid dendritic cell
PC: Principal Component
SfB: collaborative research center (Sonderforschungsbereich)
SZ: schizophrenia
TCM: central memory T cell
TEM: effector memory T cell
TNF: tumor necrosis factor
T_reg_: regulatory T cell
UMAP: uniform manifold approximation and projection

## References

1. World Health Organization. Depression and Other Common Mental Disorders: Global Health Estimates. (2017).

2. World Health Organization. Suicide worldwide in 2019: Global Health Estimates. (2021).

3. Belmaker, R. H. & Agam, G. Major Depressive Disorder. N Engl J Med 358, 55–68 (2008).

4. Goldsmith, D. R., Rapaport, M. H. & Miller, B. J. A meta-analysis of blood cytokine network alterations in psychiatric patients: comparisons between schizophrenia, bipolar disorder and depression. Mol Psychiatry 21, 1696–1709 (2016).

5. Osimo, E. F., Baxter, L. J., Lewis, G., Jones, P. B. & Khandaker, G. M. Prevalence of low-grade inflammation in depression: a systematic review and meta-analysis of CRP levels. Psychol. Med. 49, 1958–1970 (2019).

6. Benros, M. E. et al. Autoimmune Diseases and Severe Infections as Risk Factors for Mood Disorders: A Nationwide Study. JAMA Psychiatry 70, 812 (2013).

7. Dickens, C., McGowan, L., Clark-Carter, D. & Creed, F. Depression in Rheumatoid Arthritis: A Systematic Review of the Literature With Meta-Analysis: Psychosomatic Medicine 64, 52–60 (2002).

8. Müller, N. et al. The cyclooxygenase-2 inhibitor celecoxib has therapeutic effects in major depression: results of a double-blind, randomized, placebo controlled, add-on pilot study to reboxetine. Mol Psychiatry 11, 680–684 (2006).

9. Haroon, E. et al. Antidepressant treatment resistance is associated with increased inflammatory markers in patients with major depressive disorder. Psychoneuroendocrinology 95, 43–49 (2018).

10. Eisenberger, N. I. et al. Inflammation-Induced Anhedonia: Endotoxin Reduces Ventral Striatum Responses to Reward. Biological Psychiatry 68, 748–754 (2010).

11. Miller, A. H. & Raison, C. L. The role of inflammation in depression: from evolutionary imperative to modern treatment target. Nat Rev Immunol 16, 22–34 (2016).

12. Dantzer, R., O’Connor, J. C., Freund, G. G., Johnson, R. W. & Kelley, K. W. From inflammation to sickness and depression: when the immune system subjugates the brain. Nat Rev Neurosci 9, 46–56 (2008).

13. Monteiro, R. & Azevedo, I. Chronic Inflammation in Obesity and the Metabolic Syndrome. Mediators of Inflammation 2010, 1–10 (2010).

14. Raison, C. L. et al. A Randomized Controlled Trial of the Tumor Necrosis Factor Antagonist Infliximab for Treatment-Resistant Depression: The Role of Baseline Inflammatory Biomarkers. JAMA Psychiatry 70, 31 (2013).

15. Köhler-Forsberg, O., et al. Efficacy of anti-inflammatory treatment on major depressive disorder or depressive symptoms: meta-analysis of clinical trials. Acta Psychiatr Scand 139, 404–419 (2019).

16. Khandaker, G. M., Zimbron, J., Lewis, G. & Jones, P. B. Prenatal maternal infection, neurodevelopment and adult schizophrenia: a systematic review of population-based studies. Psychol Med 43, 239–257 (2013).

17. Potvin, S. et al. Inflammatory Cytokine Alterations in Schizophrenia: A Systematic Quantitative Review. Biological Psychiatry 63, 801–808 (2008).

18. Costi, S. et al. Peripheral immune cell reactivity and neural response to reward in patients with depression and anhedonia. Transl Psychiatry 11, 565 (2021).

19. Pan, W. et al. Cytokine Signaling Modulates Blood-Brain Barrier Function. CPD 17, 3729– 3740 (2011).

20. Banks, W. A., Kastin, A. J. & Gutierrez, E. G. Penetration of interleukin-6 across the murine blood-brain barrier. Neuroscience Letters 179, 53–56 (1994).

21. Gutierrez, E. G., Banks, W. A. & Kastin, A. J. Murine tumor necrosis factor alpha is transported from blood to brain in the mouse. Journal of Neuroimmunology 47, 169–176 (1993).

22. Varghese, S. M. et al. Unraveling the Role of the Blood-Brain Barrier in the Pathophysiology of Depression: Recent Advances and Future Perspectives. Mol Neurobiol (2024) doi:10.1007/s12035-024-04205-5.

23. Cathomas, F. et al. Circulating myeloid-derived MMP8 in stress susceptibility and depression. Nature 626, 1108–1115 (2024).

24. Stoeckius, M. et al. Simultaneous epitope and transcriptome measurement in single cells. Nat Methods 14, 865–868 (2017).

25. Zheng, G. X. Y. et al. Massively parallel digital transcriptional profiling of single cells. Nat Commun 8, 14049 (2017).

26. Hao, Y. et al. Dictionary learning for integrative, multimodal and scalable single-cell analysis. Nat Biotechnol 42, 293–304 (2024).

27. Hao, Y. et al. Integrated analysis of multimodal single-cell data. Cell 184, 3573–3587.e29 (2021).

28. Stuart, T. et al. Comprehensive Integration of Single-Cell Data. Cell 177, 1888–1902.e21 (2019).

29. Butler, A., Hoffman, P., Smibert, P., Papalexi, E. & Satija, R. Integrating single-cell transcriptomic data across different conditions, technologies, and species. Nat Biotechnol 36, 411–420 (2018).

30. Satija, R., Farrell, J. A., Gennert, D., Schier, A. F. & Regev, A. Spatial reconstruction of single-cell gene expression data. Nat Biotechnol 33, 495–502 (2015).

31. R Foundation for Statistical Computing. R Core Team (2021). R: A language and environment for statistical computing.

32. Choudhary, S. & Satija, R. Comparison and evaluation of statistical error models for scRNA-seq. Genome Biol 23, 27 (2022).

33. Alvarez-Mon, M. A. et al. Abnormal Distribution and Function of Circulating Monocytes and Enhanced Bacterial Translocation in Major Depressive Disorder. Front. Psychiatry 10, 812 (2019).

34. Cole, J. J. et al. No evidence for differential gene expression in major depressive disorder PBMCs, but robust evidence of elevated biological ageing. Transl Psychiatry 11, 404 (2021).

35. Li, X., Long, J., He, T., Belshaw, R. & Scott, J. Integrated genomic approaches identify major pathways and upstream regulators in late onset Alzheimer’s disease. Sci Rep 5, 12393 (2015).

36. Duan, R. et al. A De Novo Frameshift Mutation in TNFAIP3 Impairs A20 Deubiquitination Function to Cause Neuropsychiatric Systemic Lupus Erythematosus. J Clin Immunol 39, 795–804 (2019).

37. Visser, L. et al. Expression of the EGF-TM7 receptor CD97 and its ligand CD55 (DAF) in multiple sclerosis. Journal of Neuroimmunology 132, 156–163 (2002).

38. Beurel, E., Toups, M. & Nemeroff, C. B. The Bidirectional Relationship of Depression and Inflammation: Double Trouble. Neuron 107, 234–256 (2020).

39. Fusté, M. et al. Reduced expression of SP1 and SP4 transcription factors in peripheral blood mononuclear cells in first-episode psychosis. Journal of Psychiatric Research 47, 1608– 1614 (2013).

40. Pinacho, R. et al. The transcription factor SP4 is reduced in postmortem cerebellum of bipolar disorder subjects: control by depolarization and lithium. Bipolar Disorders 13, 474– 485 (2011).

41. Von Marschall, Z. et al. Effects of Interferon Alpha on Vascular Endothelial Growth Factor Gene Transcription and Tumor Angiogenesis. JNCI Journal of the National Cancer Institute 95, 437–448 (2003).

42. Belardelli, F., Ferrantini, M., Proietti, E. & Kirkwood, J. M. Interferon-alpha in tumor immunity and immunotherapy. Cytokine & Growth Factor Reviews 13, 119–134 (2002).

43. Taniguchi, T. & Takaoka, A. The interferon-α/β system in antiviral responses: a multimodal machinery of gene regulation by the IRF family of transcription factors. Current Opinion in Immunology 14, 111–116 (2002).

44. Bhat, M. Y. et al. Comprehensive network map of interferon gamma signaling. J. Cell Commun. Signal. 12, 745–751 (2018).

45. Mostafavi, S. et al. Type I interferon signaling genes in recurrent major depression: increased expression detected by whole-blood RNA sequencing. Mol Psychiatry 19, 1267– 1274 (2014).

46. Maddock, C. et al. Psychopharmacological Treatment of Depression, Anxiety, Irritability and Insomnia in Patients Receiving Interferon-α: a Prospective Case Series and a Discussion of Biological Mechanisms. J Psychopharmacol 18, 41–46 (2004).

47. Amodio, P. et al. Mood, cognition and EEG changes during interferon α (alpha-IFN) treatment for chronic hepatitis C. Journal of Affective Disorders 84, 93–98 (2005).

48. Raison, C. L. et al. Activation of Central Nervous System Inflammatory Pathways by Interferon-Alpha: Relationship to Monoamines and Depression. Biological Psychiatry 65, 296–303 (2009).

49. Maitra, M. et al. Cell type specific transcriptomic differences in depression show similar patterns between males and females but implicate distinct cell types and genes. Nat Commun 14, 2912 (2023).

50. Morrison, F. G. et al. Reduced interleukin 1A gene expression in the dorsolateral prefrontal cortex of individuals with PTSD and depression. Neuroscience Letters 692, 204–209 (2019).

51. Sun, L. et al. Peripheral Blood Mononuclear Cell Biomarkers for Major Depressive Disorder: A Transcriptomic Approach. Depression and Anxiety 2024, 1089236 (2024).

52. Sullivan, P. F., Fan, C. & Perou, C. M. Evaluating the comparability of gene expression in blood and brain. American J of Med Genetics Pt B 141B, 261–268 (2006).

53. Sun, Z. et al. Integrated Single-Cell RNA-seq and ATAC-seq Reveals Heterogeneous Differentiation of CD4+ Naive T Cell Subsets is Associated with Response to Antidepressant Treatment in Major Depressive Disorder. Adv Sci (Weinh) 11, e2308393 (2024).

54. Hammerschlag, A. R., De Leeuw, C. A., Middeldorp, C. M. & Polderman, T. J. C. Synaptic and brain-expressed gene sets relate to the shared genetic risk across five psychiatric disorders. Psychol. Med. 50, 1695–1705 (2020).

55. Jansen, R. et al. Gene expression in major depressive disorder. Mol Psychiatry 21, 339–347 (2016).

56. Ward, J. et al. The genomic basis of mood instability: identification of 46 loci in 363,705 UK Biobank participants, genetic correlation with psychiatric disorders, and association with gene expression and function. Mol Psychiatry 25, 3091–3099 (2020).

57. Mitchell, S., Vargas, J. & Hoffmann, A. Signaling via the NFΚB system. WIREs Mechanisms of Disease 8, 227–241 (2016).

58. Pace, T. W. W. et al. Increased Stress-Induced Inflammatory Responses in Male Patients With Major Depression and Increased Early Life Stress. AJP 163, 1630–1633 (2006).

59. Jiang, Y. et al. Gypenoside-14 Reduces Depression via Downregulation of the Nuclear Factor Kappa B (NF-kB) Signaling Pathway on the Lipopolysaccharide (LPS)-Induced Depression Model. Pharmaceuticals (Basel) 16, 1152 (2023).

