## Supplementary figures for "Monocyte Dysregulation Defines an MDD-Specific Transcriptional Signature Closely Linked to Clinical MDD Traits"

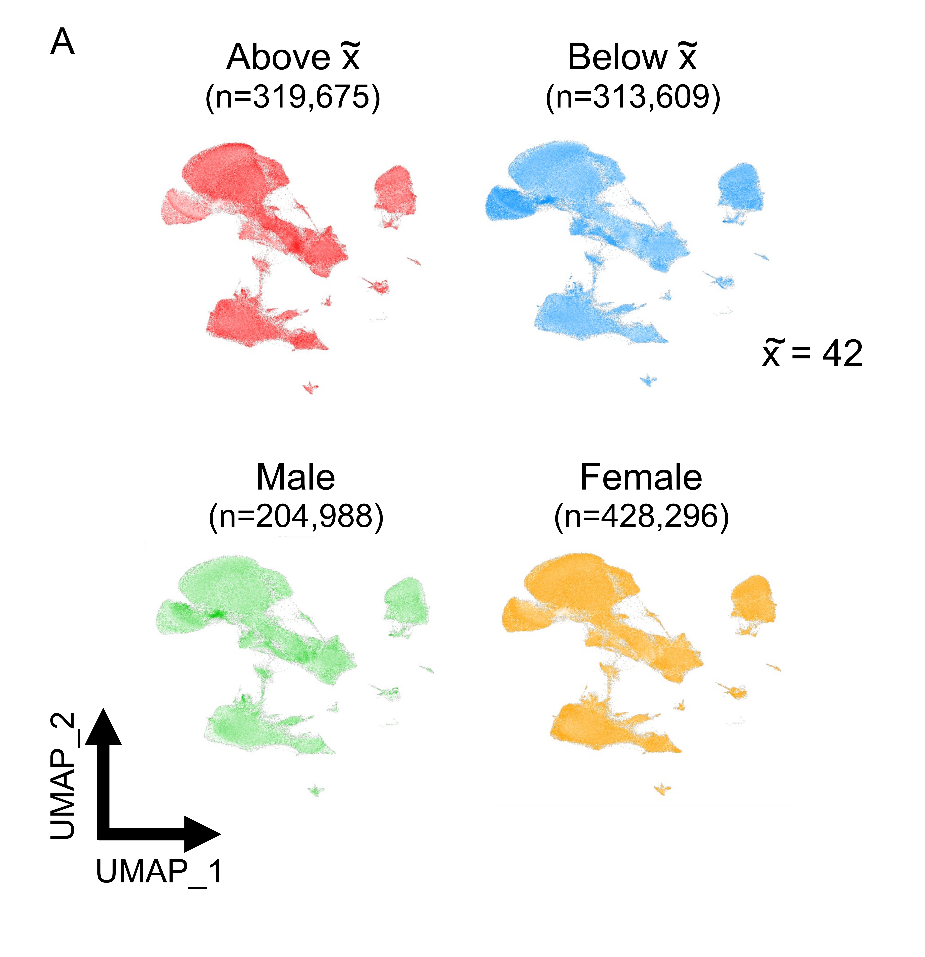

**Supplementary figure 1: Distribution of major cell types across age and sex.** No significant overrepresentation of major cell types was identified among different age and sex groups.

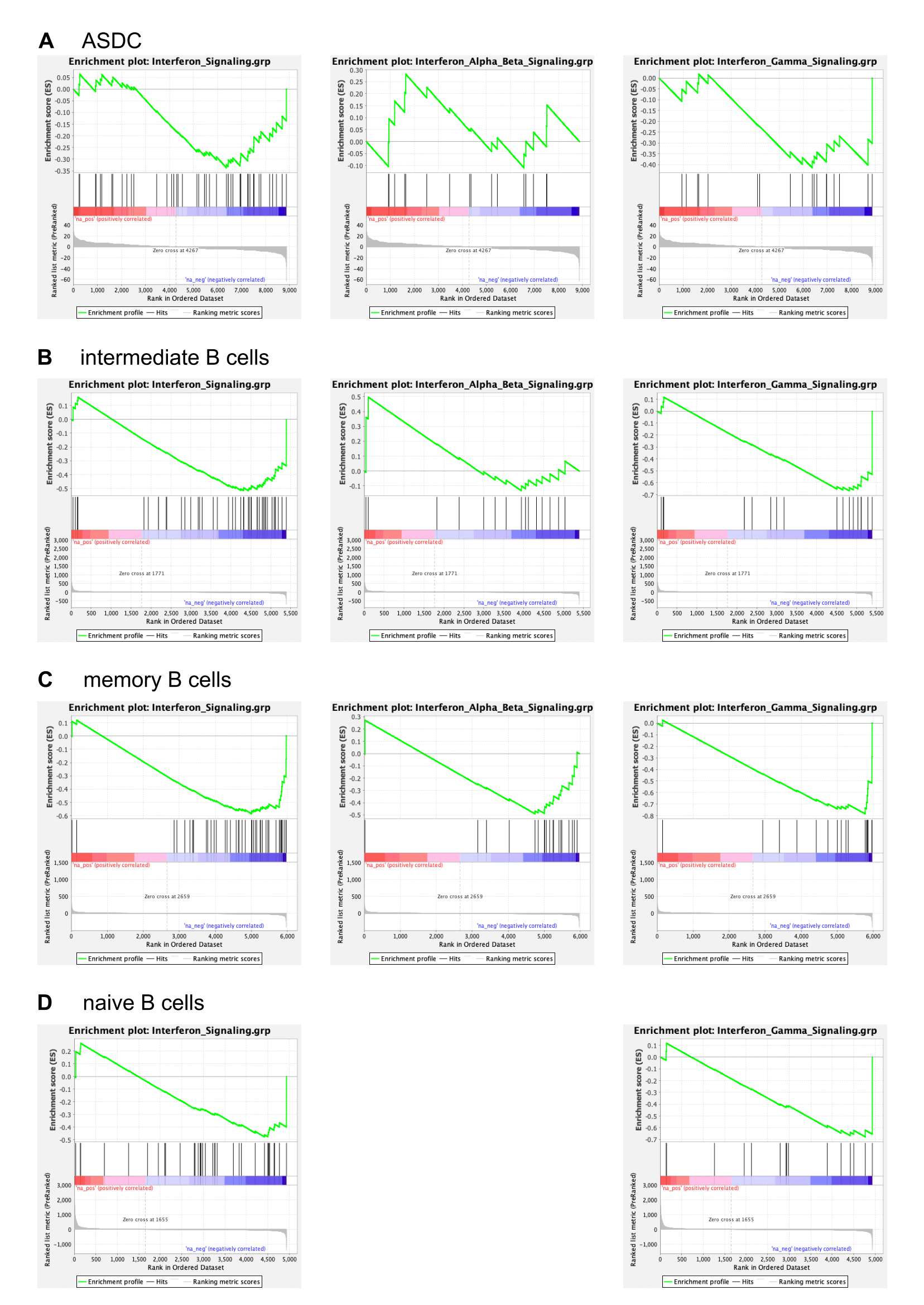

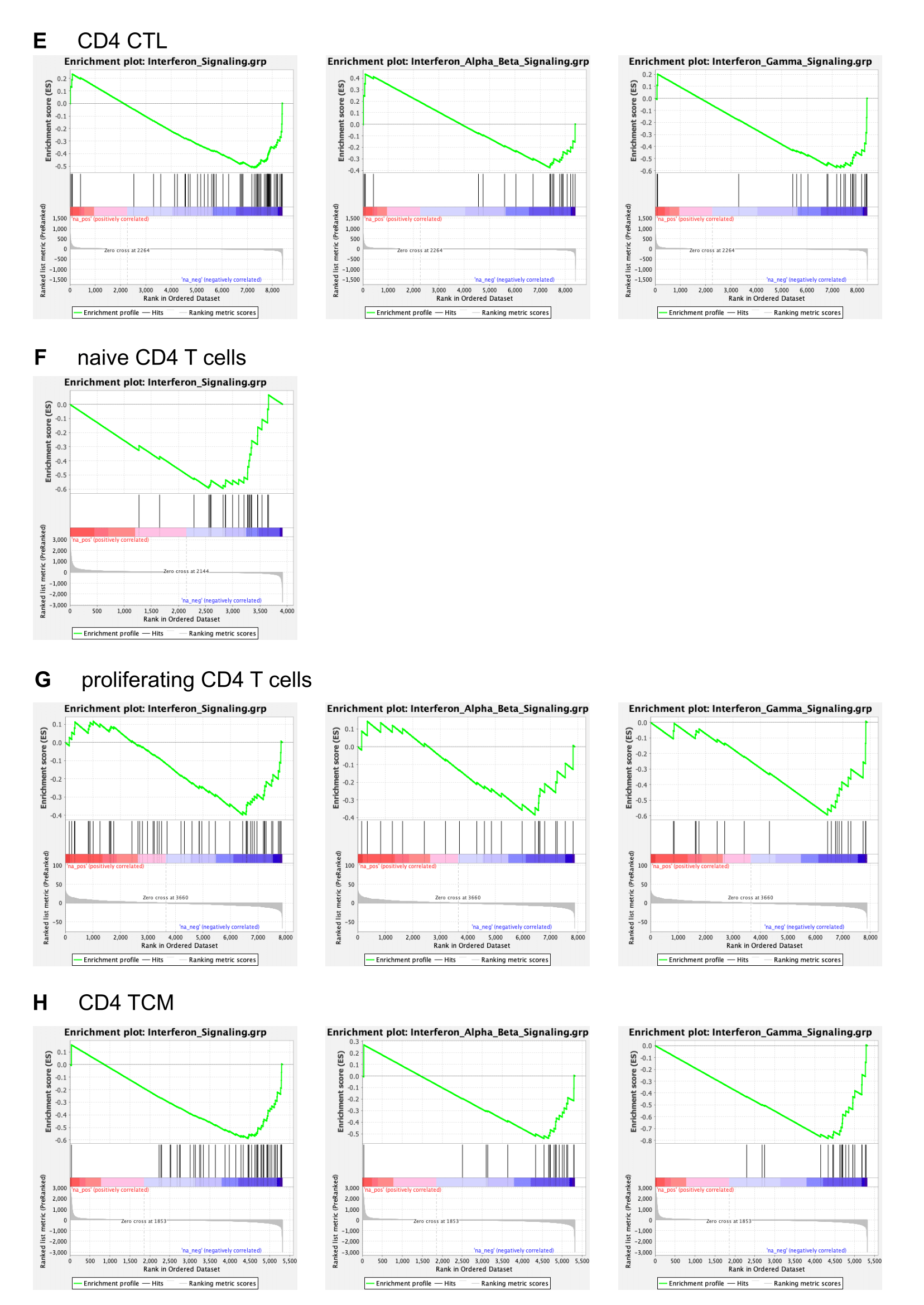

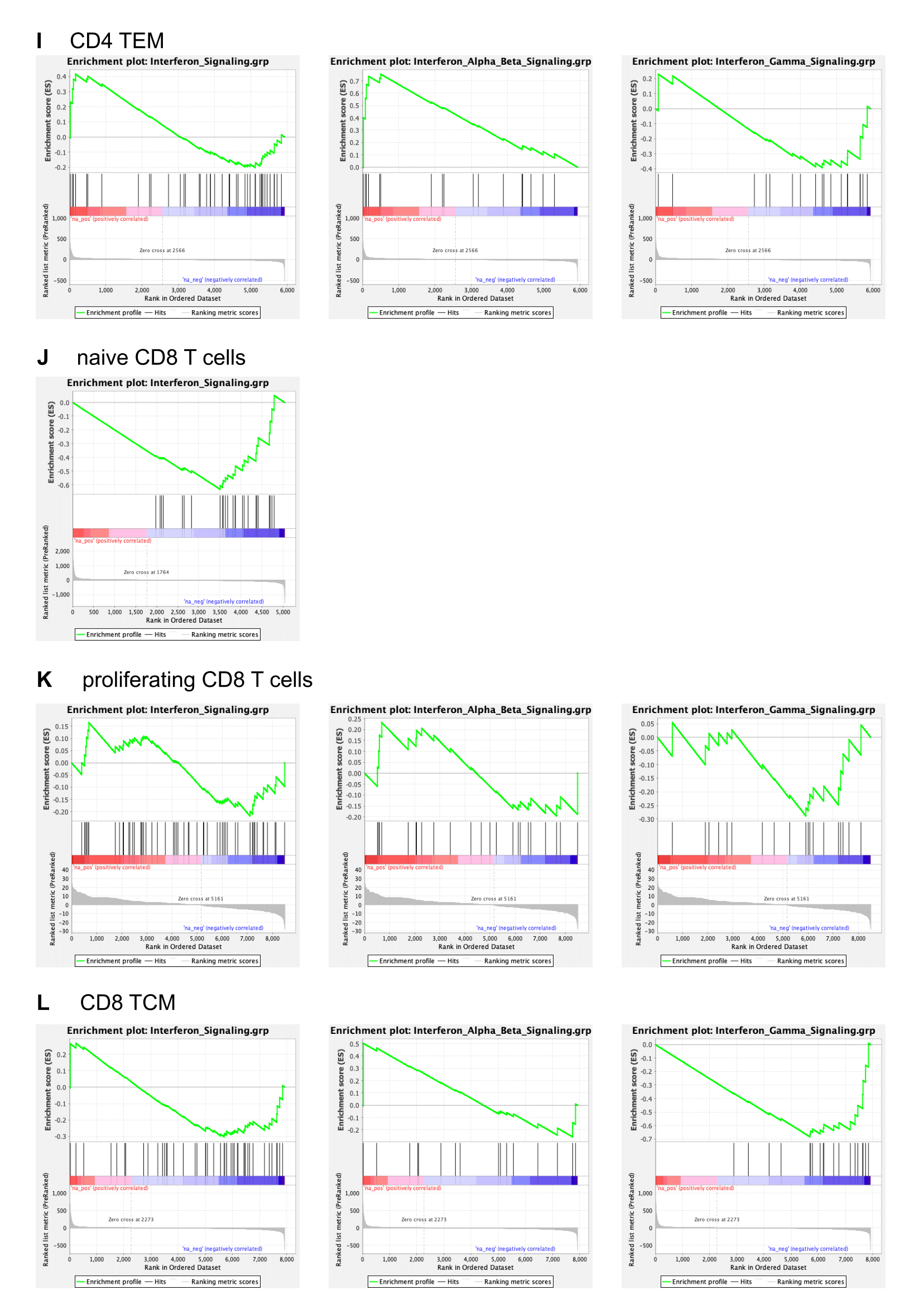

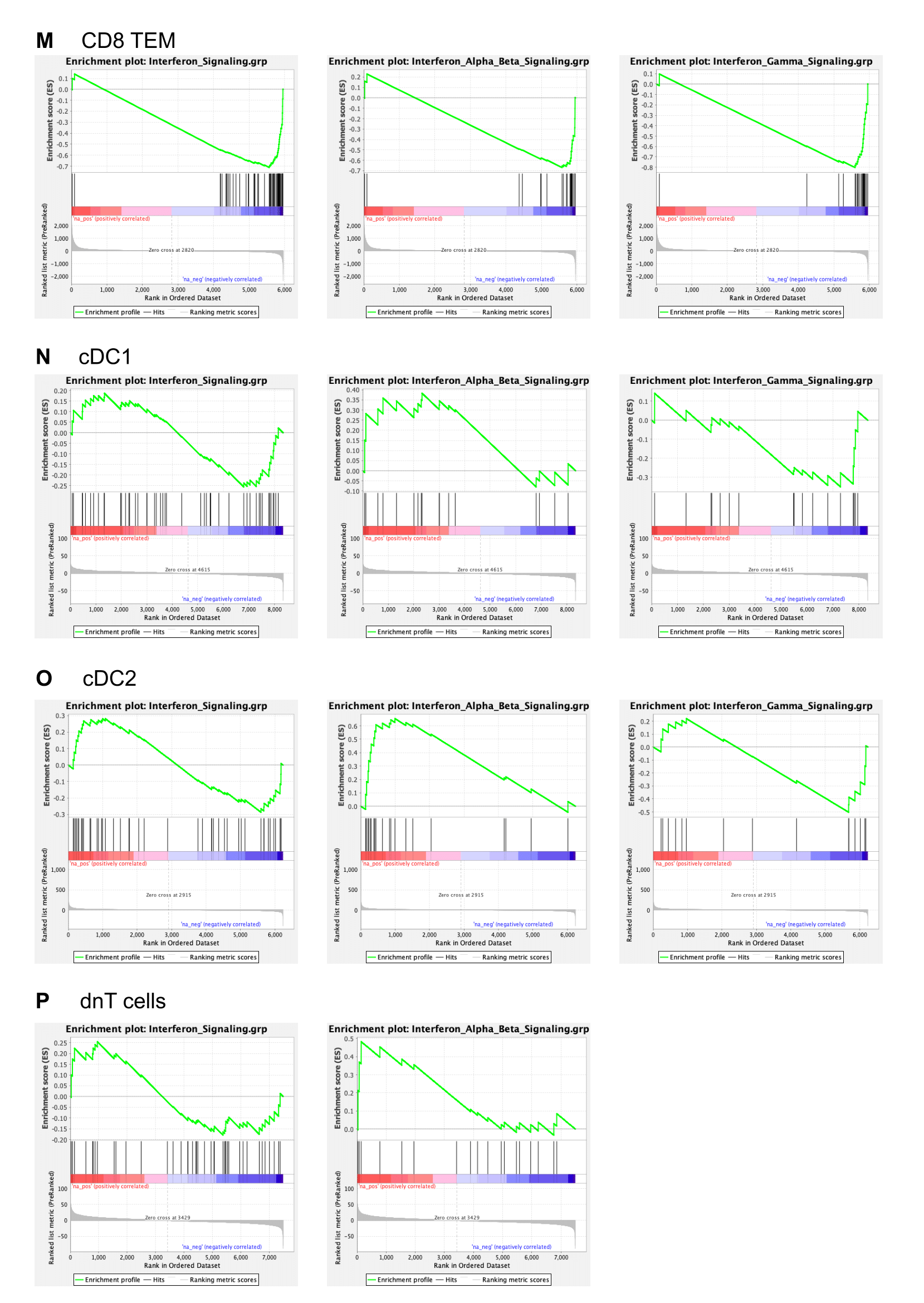

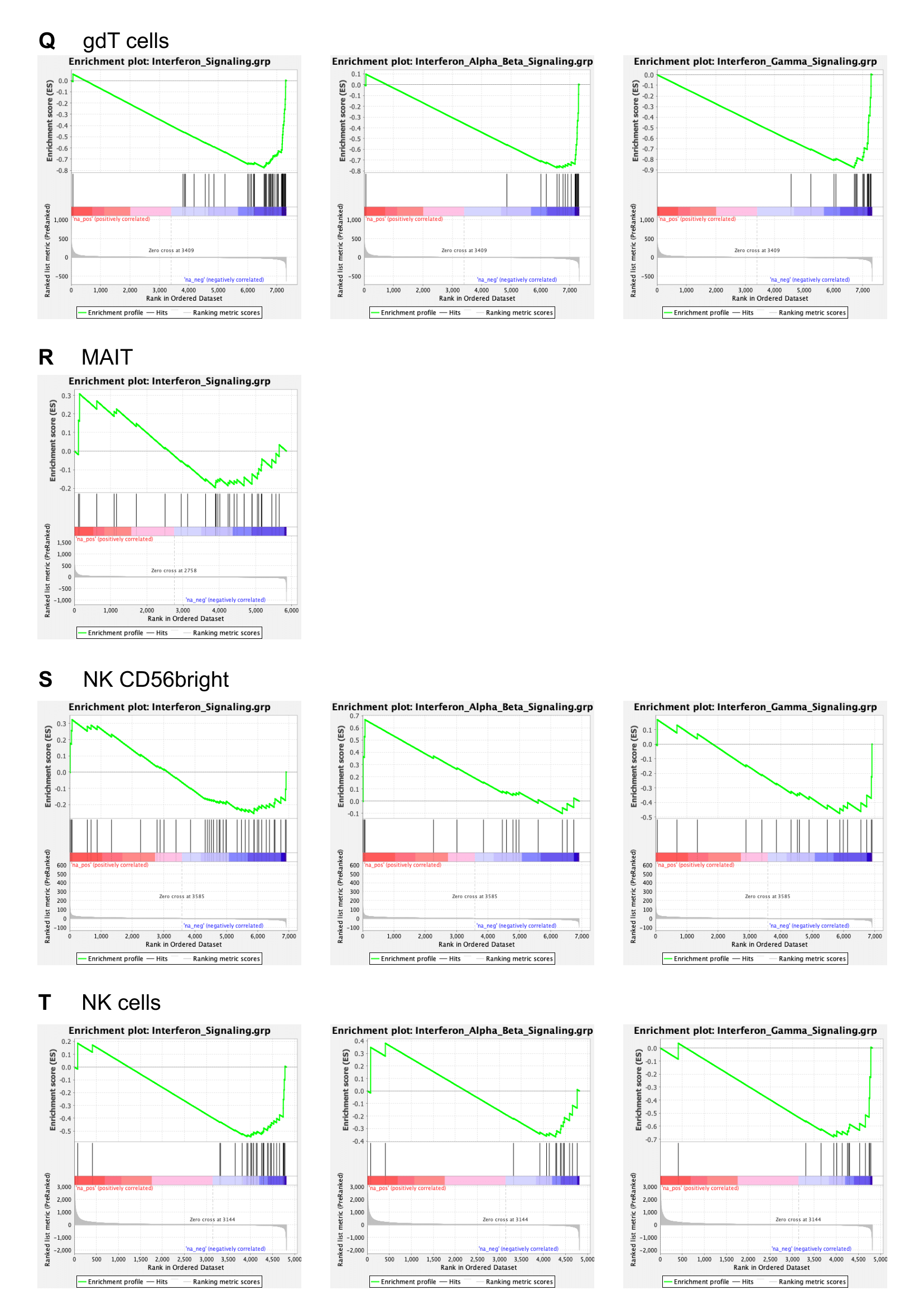

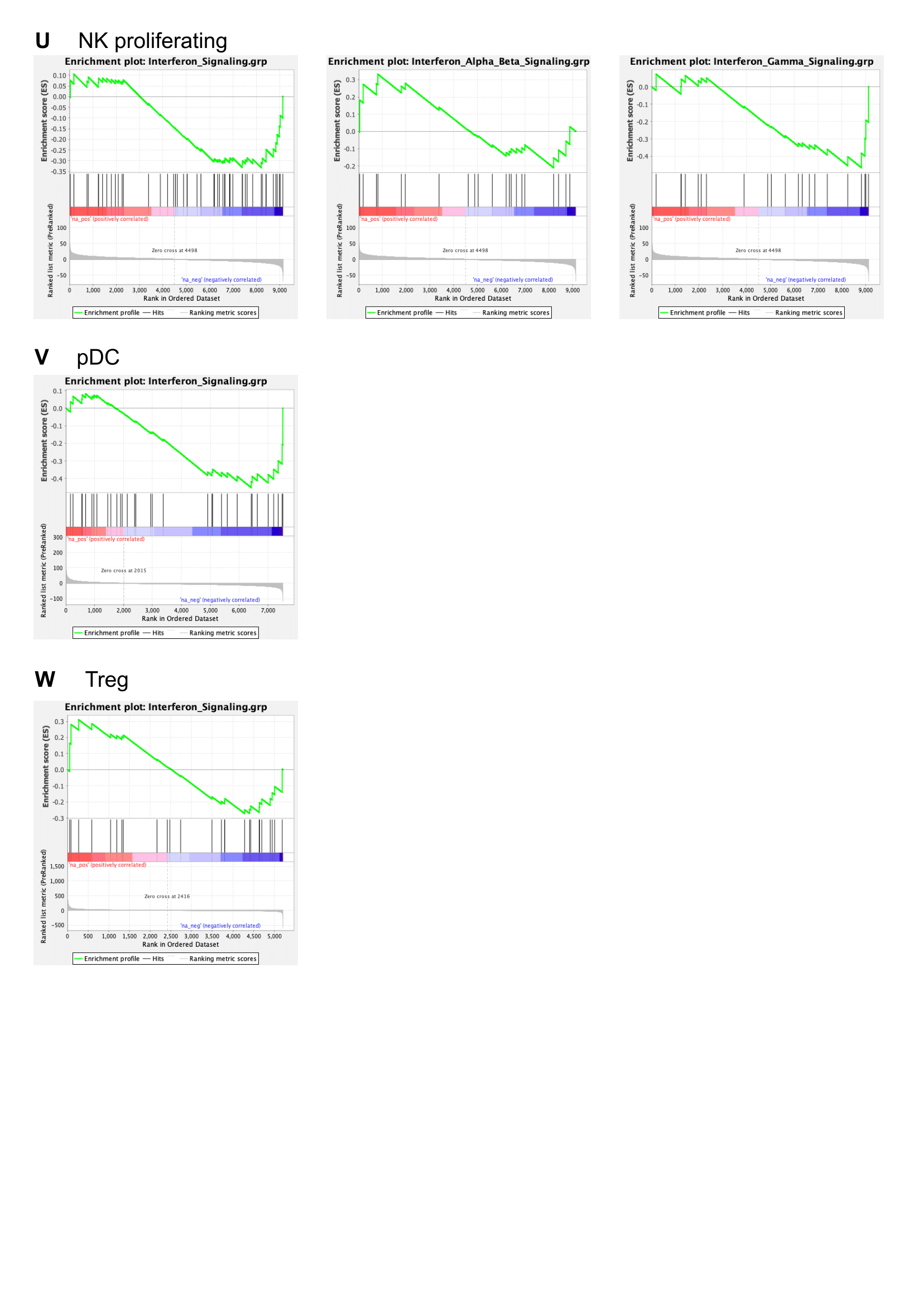

**Supplementary figure 2: Enrichment plots of interferon signaling for PBMC subtypes.**

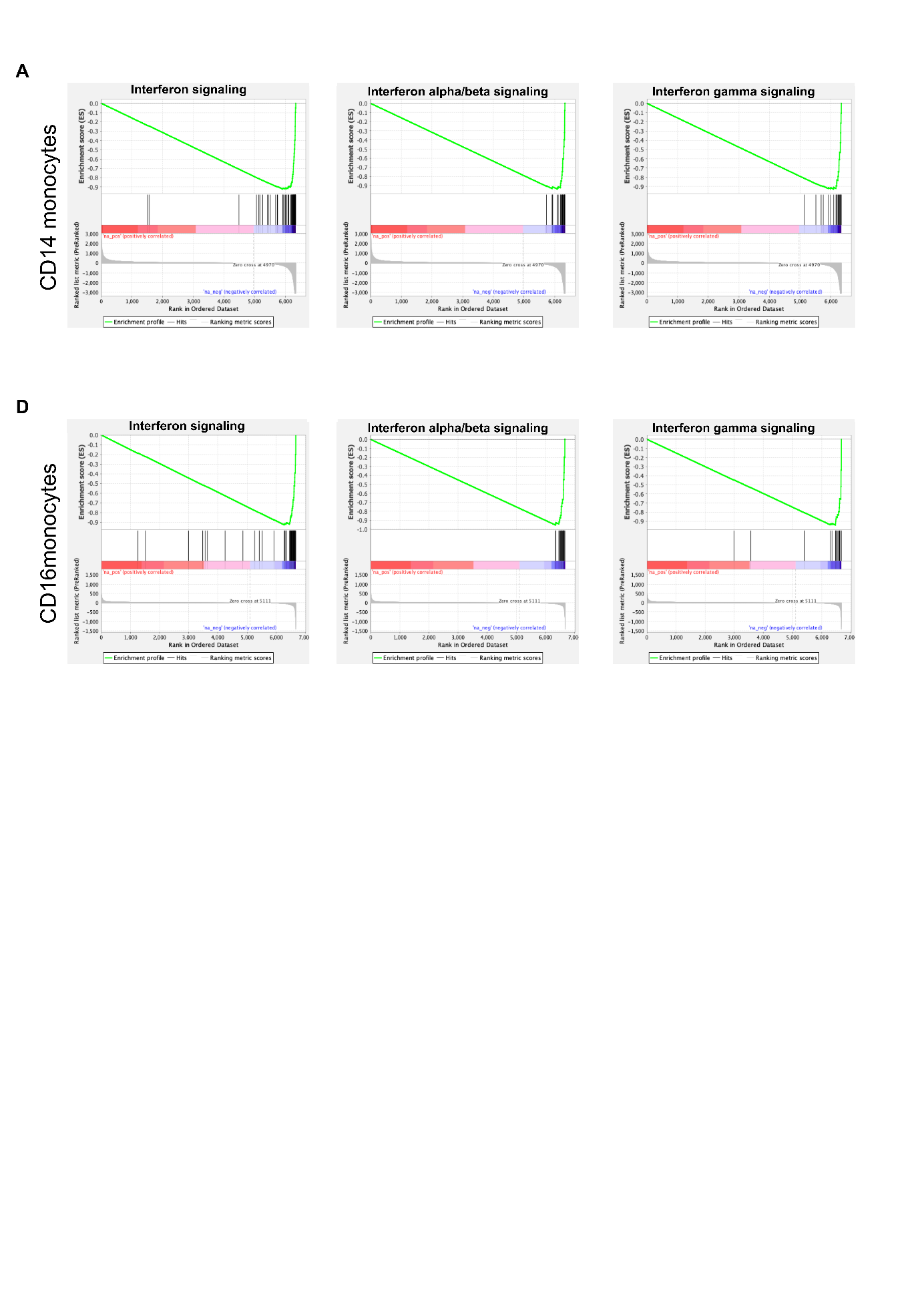

**Supplementary figure 3: Enrichment plots of interferon signaling for CD14 and CD16 monocytes.**

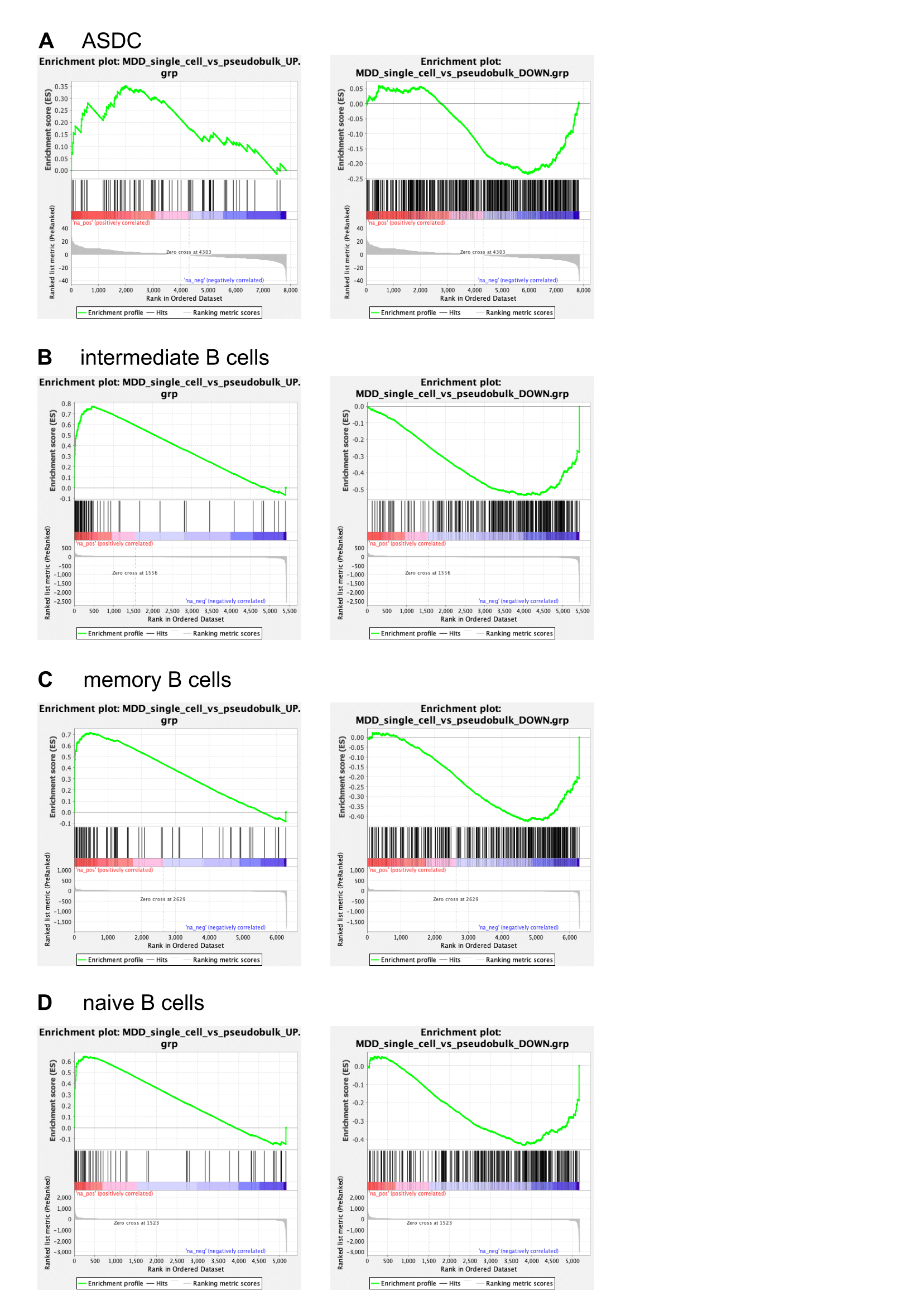

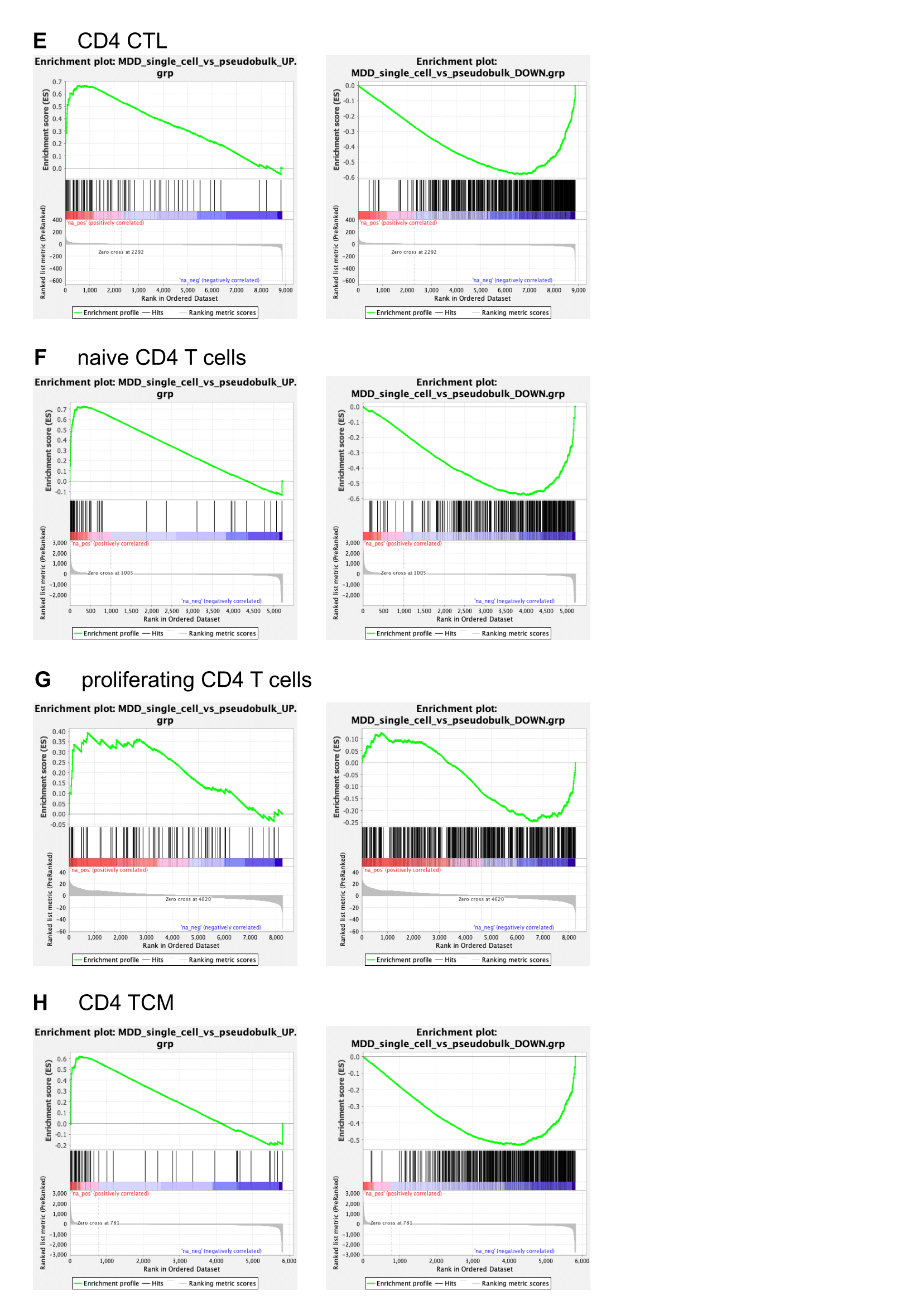

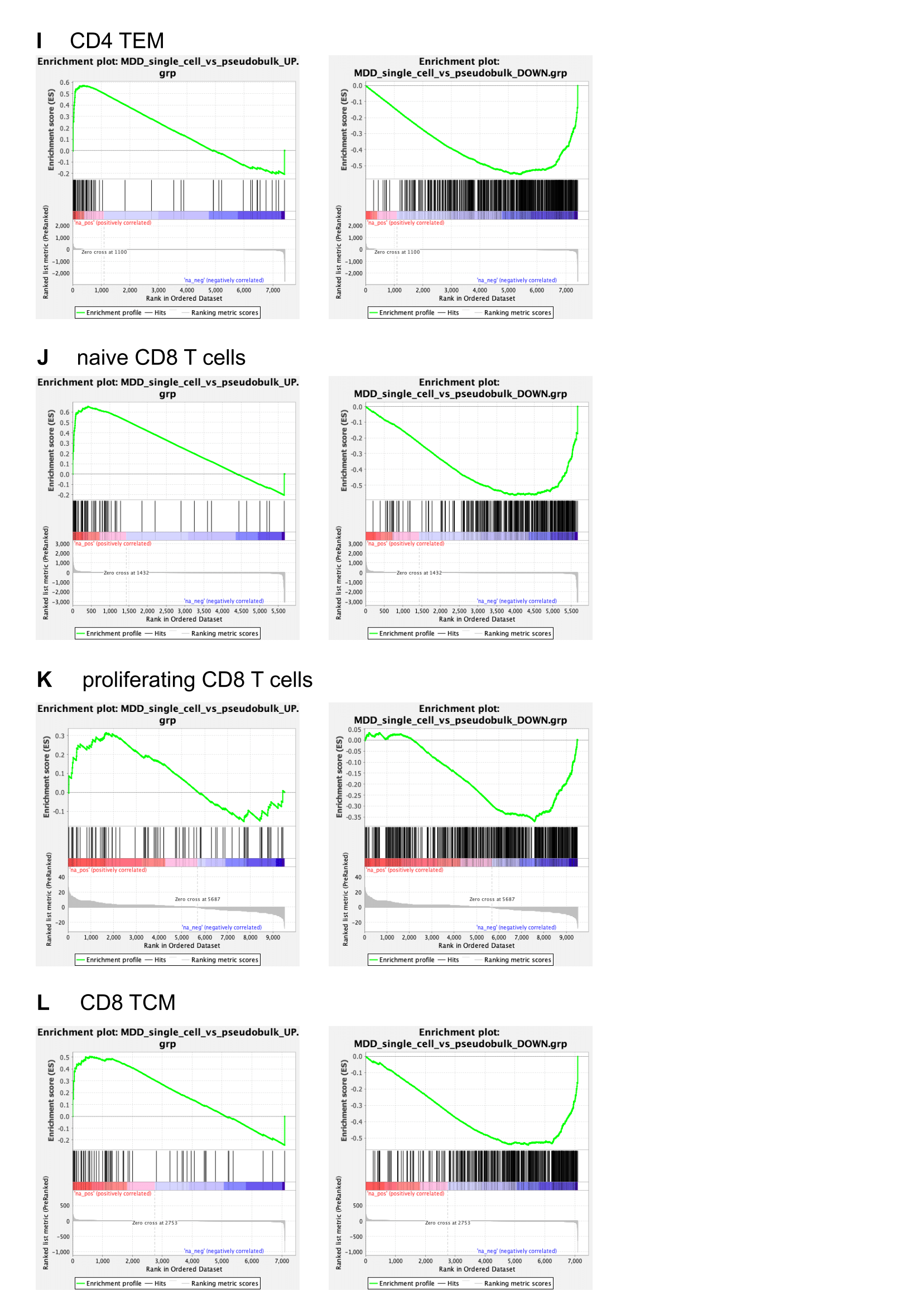

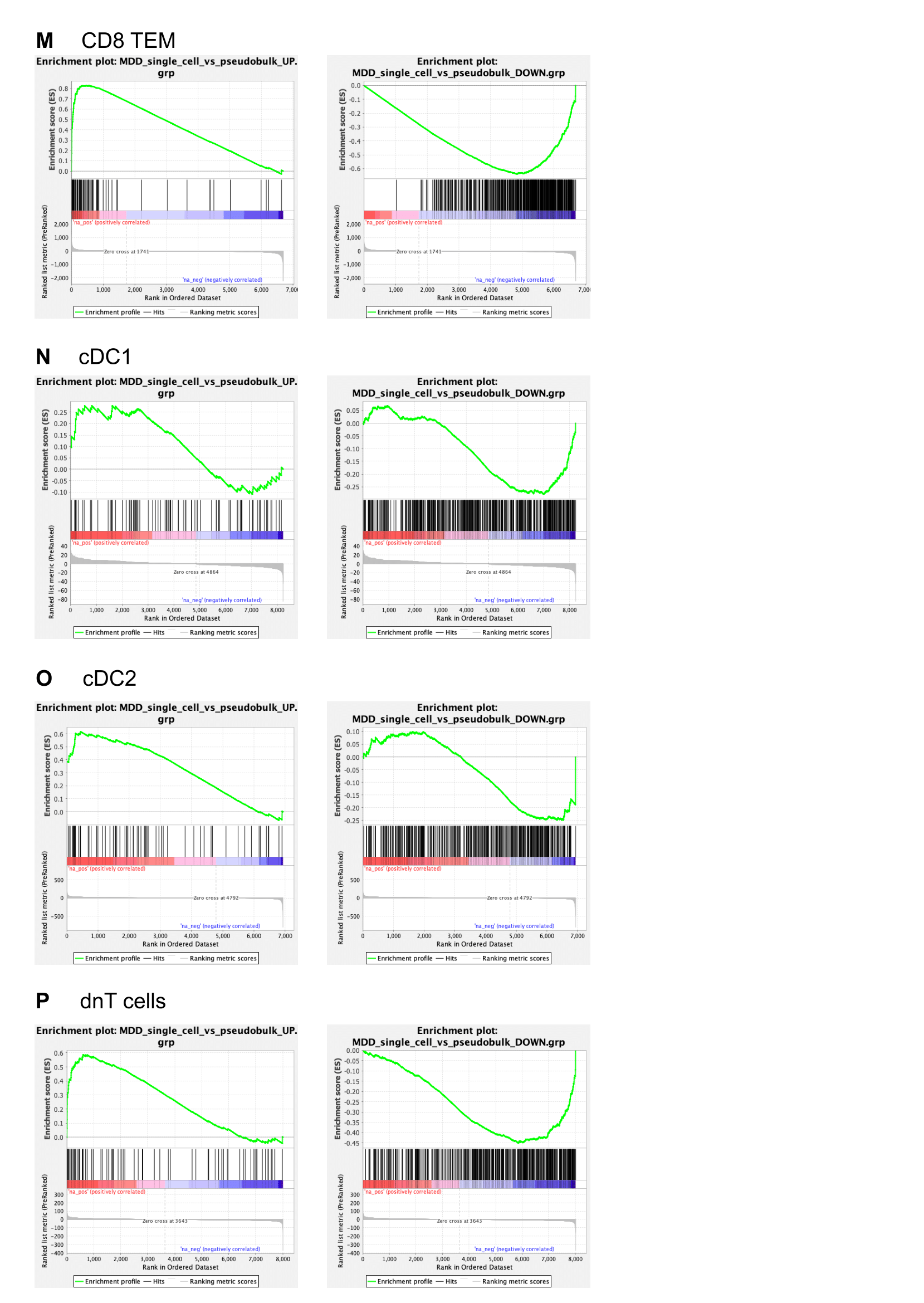

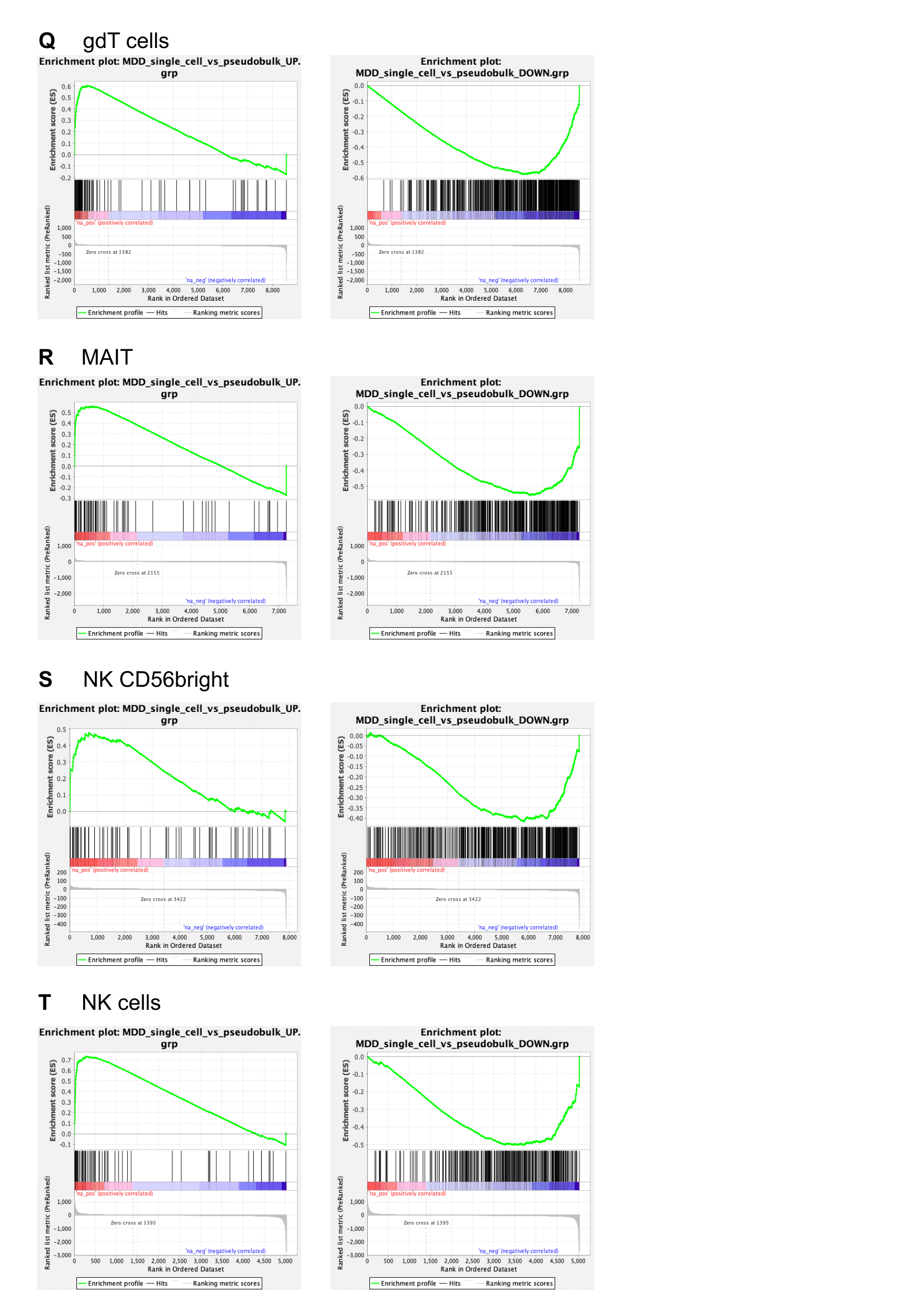

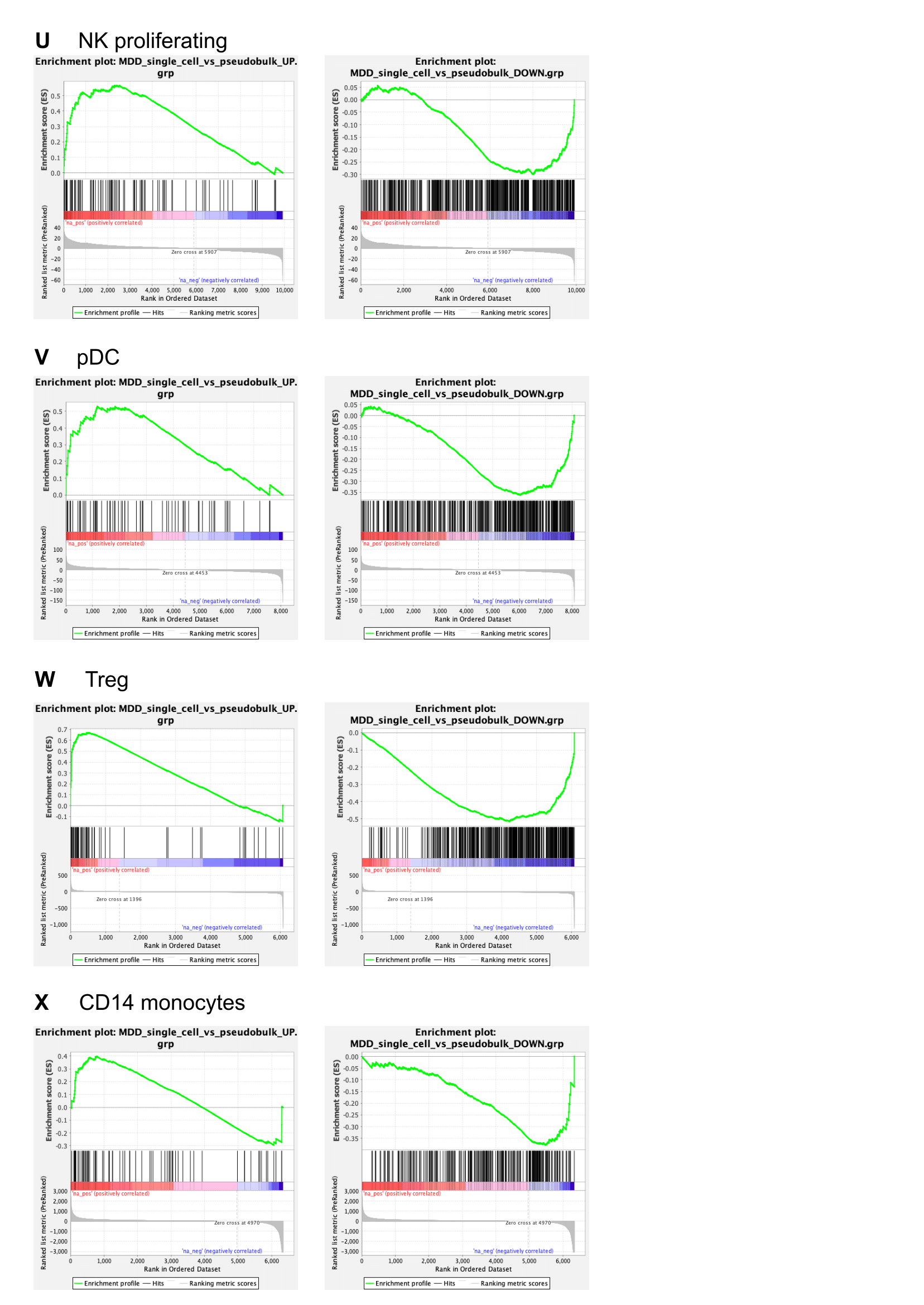

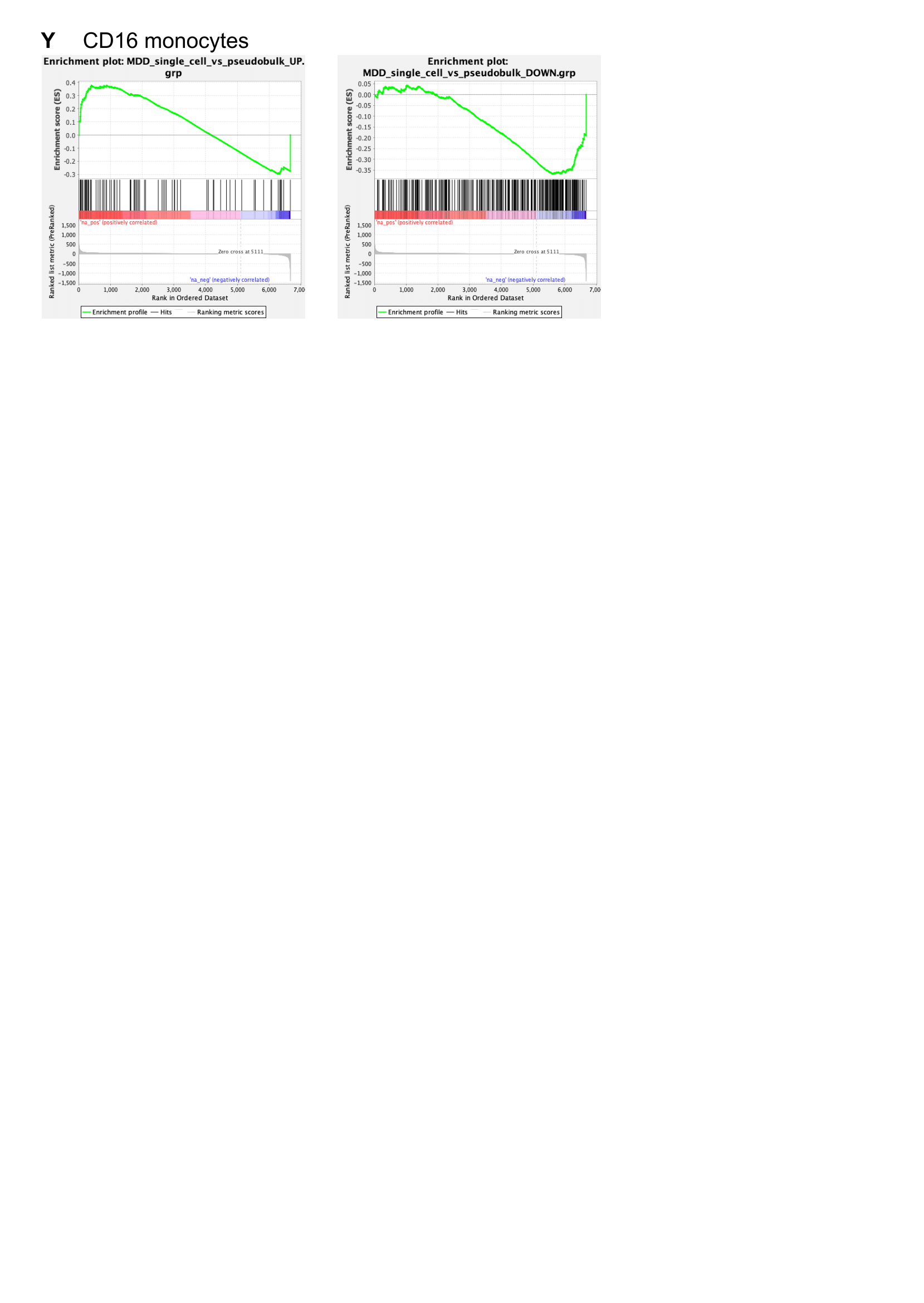

**Supplementary figure 4: Enrichment plots of the disease-specific transcriptional MDD core signature in all cellular subtypes.**
